# Paralemmin-2 is a membrane-anchored cytoskeletal constituent of Axon Initial Segments and nodes of Ranvier

**DOI:** 10.64898/2026.09.03.749080

**Authors:** Victor Macarrón-Palacios, Nicole G. Metzendorf, Greta Hultqvist, Claudio Acuna, Simon Kneilmann, Jasmine Hubrich, Liane Wüstefeld, Henrik Martens, Manfred W. Kilimann, Elisa D’Este

## Abstract

The axon initial segment (AIS) and nodes of Ranvier (NoR) are essential for action potential initiation and propagation. They share many features of their molecular architecture, and their assembly mechanisms converge on the membrane-associated periodic skeleton (MPS). Here, we identify Paralemmin-2 (Palm2) as a component of both the AIS and NoR. Palm2 depletion shortens the AIS and reduces neuronal excitability. Endogenous Palm2 in the AIS is non-periodic, but upon overexpression it associates with the MPS, localizing to actin rings and reducing βIV-spectrin abundance and periodicity. In NoR of the central and peripheral nervous system, Palm2 localizes to different subdomains - nodal or paranodal, respectively. Palm2, and its homolog Palm1, bind the deubiquitinase USP7, implicating paralemmins in proteostasis at the MPS. The complementary localizations of Palm2 and Palm1 at the AIS/NoR or axon shafts, respectively, parallel the distributions of β-spectrin and ankyrin isoforms between these axonal compartments. We propose that Palm2 modulates the submembrane cytoskeleton and its membrane attachment, and thus contributes to the assembly, functioning and remodeling of the AIS and NoR.

## Introduction

The AIS is a specialized structure located at the first ∼30 µm of the axon and responsible for the integration of synaptic inputs and action potential (AP) generation (*1, 2*). It exhibits a very dense submembrane scaffold, acts as a diffusion barrier for membrane proteins and lipids, and as a filter for intracellular traffic discriminating between axon-and dendrite-destined molecular cargo. During AIS assembly, ankyrinG (ankG) plays a pivotal role: it is one of the earliest proteins to be detected in this compartment and is required for the clustering of many AIS components (*3*). As ankG interacts with multiple proteins, including the spectrin isoform βIV-spectrin, it serves as a master scaffolding protein for building macromolecular complexes along the AIS (*4–10*). The molecular architecture of Nodes of Ranvier (NoR), the sites where the AP is regenerated along myelinated axons, shares similarities with the AIS, including a pivotal role of ankG and the presence of βIV-spectrin. However, while AIS formation is spontaneous and cell-autonomous, NoR assembly requires the interaction with myelinating glia cells (*11*).

The cytoskeleton is crucial for axon specification and AIS assembly. Particularly along the axons, the submembrane cytoskeleton assumes a highly-ordered 190 nm periodic organization, known as the Membrane-associated Periodic Skeleton (MPS) (*12–14*). This actin-spectrin lattice provides mechanical support, controls intracellular transport, endocytosis, and organizes signaling proteins and voltage-gated channels (*12, 15–22*). To orchestrate and confine these functions to specific axonal compartments, the molecular composition of the MPS differs in the AIS from the middle-distal axon (*1–3, 23*). During axon specification and AIS formation, different isoforms of MPS core proteins compartmentalize along the axon: ankG and βIV-spectrin concentrate at the AIS, whereas ankB and βII-spectrin are depleted from the AIS and populate mainly the middle and distal axons. The local concentrations of both ankG and βIV-spectrin in the AIS increase during neuronal development, although their periodic organization sets in at different time points. AnkG assumes a periodic organisation early on, within the first week of neuronal development in culture, while βIV-spectrin exhibits a clear periodicity from approximately day in vitro (DIV) 12 (*17, 24, 25*). The AIS is a dynamic structure whose length, position, and composition are finely tuned in response to changes in activity levels (reviewed in (*3, 26*)). Activity-dependent remodeling involves the ubiquitin-proteasome system, which targets ankG (*27*). While these are important insights, much remains to be learned about the AIS assembly, homeostasis, and plasticity. Which molecular mechanisms underlie the diffusion barrier and transport filter functions of the AIS? How is the developmental assembly of this huge, highly integrated molecular machine orchestrated? Probably, only a minority of AIS component proteins have been identified (*28–30*), let alone mechanistically understood.

In this study, we identify Paralemmin-2 (Palm2) as a novel constituent and putative regulator of the AIS and NoR submembrane cytoskeleton. Palm2 is a member of the Paralemmin protein family, together with the isoforms Palm1, Palm3, and Palmdelphin (Palmd) (*31*). They share a molecular architecture consisting of an N-terminal coiled-coil domain, which is thought to mediate interaction with the actin cytoskeleton, and a C-terminal CaaX lipidation motif, which is responsible for anchoring to the plasma membrane. The intervening sequences, meanwhile, are more divergent and are predicted to be largely intrinsically unstructured. Palm1, the founding member and most abundant isoform of this family, was recently shown to bind the N-terminal region of βII-spectrin, localize to the MPS at the actin-spectrin junction complex, and enhance the nanoscale periodicity of the MPS (*32*). Current knowledge suggests that Palm1 is attached to the plasma membrane through its C-terminal lipid anchor, and reaches into the cytoplasm with its N-terminal parts, stabilizing the MPS and tightening its association with the membrane. Furthermore, overexpression of Palm1 induces filopodia formation and spine maturation, while knock-down has the opposite effect (*33*). Palm3 is essential for the stability of the cortical lattice of auditory hair cells – a highly regular submembrane actin-spectrin skeleton similar to the MPS - as well as for the spatial organization of intrinsic proteins of the lateral plasma membrane to which Palm3 connects the cortical lattice (*34*). Regarding Palmd, its membrane-anchored splice variant promotes the formation of cell processes by neocortical neural progenitor cells through interaction with adducin, an actin capping protein and MPS component (*35*), whereas the cytoplasmic Palmd splice variant enhances the resilience of the perinuclear actin cap against shear stress in endothelial cells (*36*), promotes myoblast differentiation and muscle regeneration (*37*), and is a constituent of the Z-discs of cardiomyocytes (*38*). As for medical involvements, Palm3 deficiency causes hearing impairment (*34*), Palmd is implicated in calcific aortic valve stenosis and arterial calcification (*39–41*), and Palm1 has been linked to migraine (*42*) and maternal gestational diabetes mellitus (*43*). Collectively, these findings show that paralemmins bind, organize, and strengthen various actin-based cytoskeletal structures, often in conjunction with spectrin and with the plasma membrane. Thus, paralemmins seem to constitute a novel molecular principle in the regulation of the actin cytoskeleton, but an understanding of their biological roles and their molecular mechanisms is still only beginning to emerge.

For Palm2, knowledge of its expression pattern, subcellular localizations and biological contexts has been particularly scarce. In the present study, we explored the role of Palm2 in the nervous system. Being the closest relative of Palm1 by amino acid (aa) sequence and overall molecular architecture, we hypothesized a similar role for Palm2, perhaps in other subcellular compartments. By immunofluorescence (IF) of brain sections and cultured neurons, we noted an enrichment of Palm2 at the AIS and NoR but, surprisingly, in different compartments of the NoR in the central (CNS) and peripheral (PNS) nervous systems. Ablation or overexpression of Palm2 impact the protein composition of the AIS and electrophysiological properties of the neurons. Overall, Palm1 and Palm2 display properties that are in part similar, in part different or even opposite, suggesting complementary roles in the regulation of the submembrane actin-spectrin cytoskeleton in different compartments of the neuronal plasma membrane.

## Results

### Palm2 is widely expressed in the nervous system and concentrates at the AIS and NoR

We began our investigation by identifying the tissues in which Palm2 is expressed. An immunoblot survey of mouse organs detected Palm2-CaaX only in the brain and eye (Fig. 1A). This “standard” form of murine Palm2 ends with a C-terminal CaaX motif to which a lipid membrane anchor is posttranslationally attached, is 377 amino acids (aa) long, but it migrates at ∼72 kDa because it is highly acidic (*44*). Additional variants of Palm2 are generated by differential splicing and alternative promoter choice (*44*) (see Discussion), of which a band of ∼250 kDa, probably representing the splice variant Palm2-AKAP2, was detected in adult lung, uterus, heart and, very faintly, eye. Samples from newborn mouse heads and bodies both gave marked 250 kDa bands, while the 72 kDa band was only detected in newborn head. Quantitative immunoblot analysis, calibrated with bacterially expressed recombinant paralemmins, showed that Palm1 is the predominant isoform in adult mouse brain (0.2% of total brain protein), whereas Palm2 and the other isoforms are by two orders of magnitude less abundant (Palm2: 0.007%; Palm3: 0.002%; Palmd: 0.002%, see methods).

**Figure 1:**
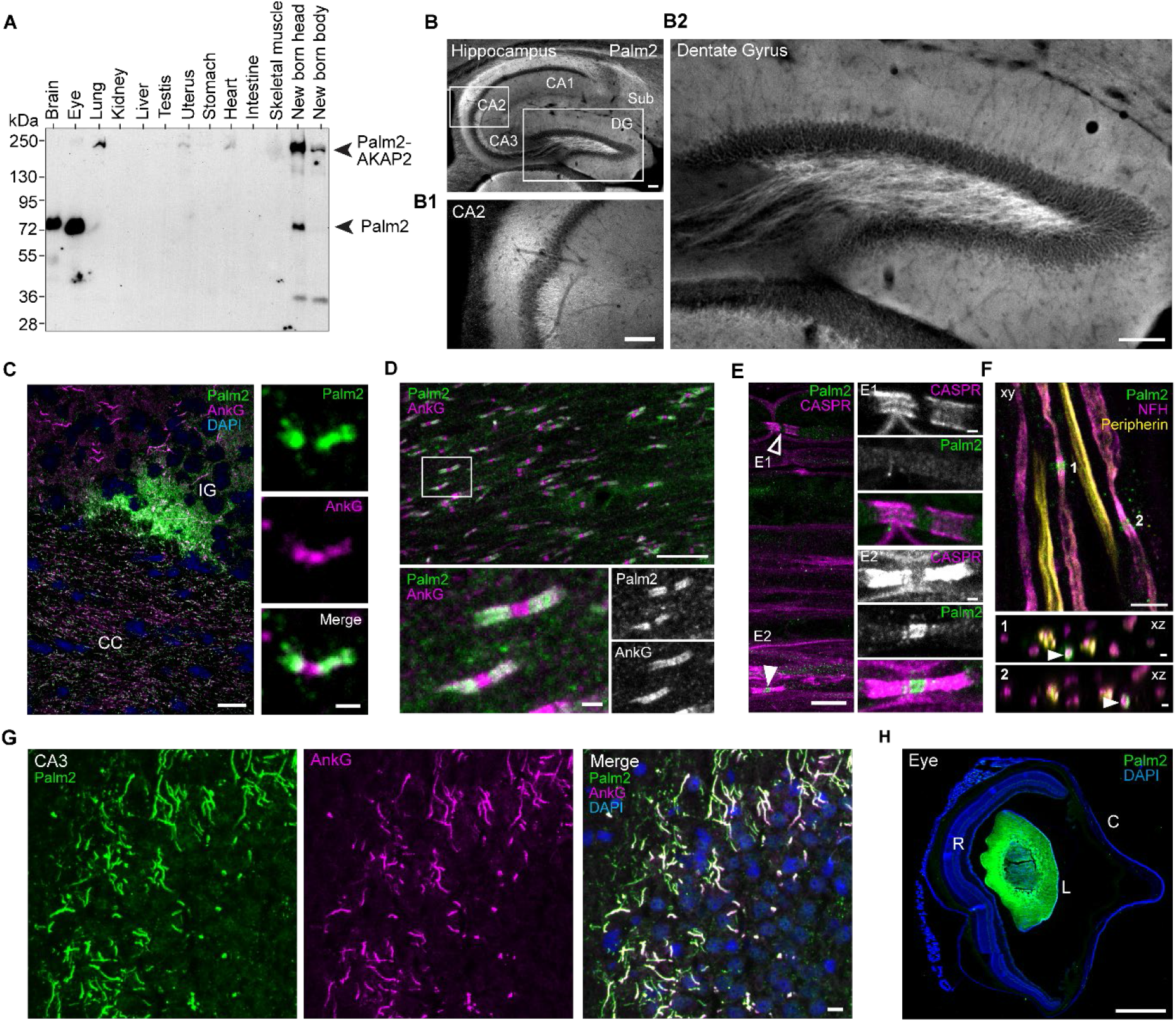
Palm2 expression and localization in mouse organs. (A) Immunoblot of Palm2 expression in the indicated mouse organs (p21 antibody). Bands at ∼72 kDa and ∼250 kDa correspond to Palm2-CaaX and Palm2-AKAP2, respectively. Uncropped blot is shown in file S2. (B) Palm2 (p22 antibody) in the hippocampal formation (CA1-3, dentate gyrus [DG], subiculum [Sub]). Scale bar: 100 µm. (B1) Palm2-positive neuropil layers in the CA2 area. Scale bar: 50 µm. (B2) Close-up image of the dentate gyrus shows Palm2-positive mossy-fiber axons emanating from the granule cell layer. Scale bar: 100 µm. (C) Palm2 (p22 antibody) is present in the indusium griseum (IG) and at the NoR paranodes of the corpus callosum (CC), identified by ankG. Right: Close-up images of a NoR in the CC. Scale bars: 10 µm on the left, 1 µm on the right. (D) Palm2 (1A8 antibody) in the cerebellar medulla. The ankG antibody recognizes both nodal gaps (ankG-480 kDa in axons) and paranodes (ankG-190 kDa in oligodendrocytes). White box indicates close-up merged and single-channel images at the bottom. Scale bars: 10 µm in upper panel, 1 µm in close-ups. (E) Palm2 (p22 antibody) in teased sciatic nerve fibers. CASPR identifies paranodal regions. (E1) Empty and (E2) solid arrowheads point at nodal gaps that are Palm2-negative and -positive, respectively. On the right, close-up merged and single channel images of the two nodes indicated on the left. Brightness is adjusted to the same level for all panels. Scale bars: 10 µm in upper panel, 1 µm in close-ups. (F) Teased sciatic nerve fibers stained against Palm2 (p22 antibody), neurofilament-H (NFH) as marker of sensory neurons, and peripherin as marker of motor neurons. Top: xy scan (z-stack projection), Bottom: xz scans (y-stack projection) of the nodal gaps, as indicated by numbers in the xy image. Arrowheads point at the Palm2-positive NoR. Scale bars: 5 µm in upper panel, 1 µm in xz scans. (G) Palm2 (p21 antibody) in the AIS of the CA3 region of the hippocampus, colocalizing with ankG. Scale bar: 20 µm. (H) Medio-sagittal section through a mouse eye shows Palm2 staining (p21 antibody) only in the lens (L) but not in the retina (R) or cornea (C). Scale bar: 500 µm. See Table S1 for experimental details.

We then surveyed sections of mouse brain by IF and observed Palm2 in many areas, prominently in the hippocampal formation (Fig. 1B; Fig. S1A). Finely dispersed Palm2 immunoreactivity was seen in neuropil-rich strata, most intensely in the CA2 stratum oriens (Fig.1B1), and some tracts of unmyelinated axons were standing out, such as mossy fibers or axons projecting from the CA1 to the subiculum (Fig. 1B,B2; Fig. S1A). Other prominently Palm2-positive regions of the brain were the indusium griseum (Fig. 1C; Fig. S1B) and the fasciola cinerea, both also belonging to the hippocampal formation, and the lining of ventricles (Fig. S1C).

In myelin-rich structures such as the corpus callosum (Fig. 1C; Fig. S1B,C), the cerebellar medulla (Fig. 1D) or myelin bundles in the striatum (Fig. S1C), we noted coarsely punctate Palm2 immunostaining. High magnification and double-IF with anti-ankG revealed a staining pattern typical for the NoR, with ankG labeling the nodal gap and, less intensely, the flanking paranodes, while Palm2 stained only the paranodes (Fig. 1C-D; Fig. S1D-E). This paranodal localization of Palm2 was consistently observed in several brain regions (corpus callosum, cerebellum, striatum, brainstem). In contrast, at the NoR of sciatic nerves, *i.e.* in the PNS, we found Palm2 to concentrate at the nodal gaps, but not at the Caspr-positive paranodes, and only in some fibers (Fig. 1E). Fibers containing Palm2-positive NoR were enriched in neurofilament-H, a marker for sensory fibers, rather than peripherin, a marker of motor neurons (Fig. 1F). Vertical optical sectioning through the nodal gap showed that Palm2 forms a ring surrounding NFH, suggesting its positioning at the axonal plasma membrane.

Palm2 IF of primary hippocampal neurons in culture prominently labeled the AIS under various fixation/permeabilization methods (see the following chapters). However, in tissue sections after conventional PFA fixation, Triton X-100 permeabilization and heat-treatment for antigen retrieval (e.g. Figs. 1B, S1A), AIS labeling for Palm2 was very faint at best. In contrast, fresh-frozen and acetone-postfixed brain sections displayed AIS staining for Palm2 in the hippocampal CA1-3 pyramidal layers (Figs. 1G, S1F), in the striatum and in the neocortex, but none in the cerebellum or retina, suggesting different concentrations or accessibilities of Palm2 in different AIS populations. AIS staining for Palm2 was also observed in guinea-pig neocortex (Fig. S1G; see Discussion regarding the expression of Palm2 splice variants in different animal species).

Of all organs analyzed by immunoblotting, Palm2 was most abundant in the whole eye (Fig. 1A). IF showed that Palm2 was exclusively concentrated in the lens (Fig. 1H), whereas no specific staining was seen in the retina even at high magnification and with various fixation conditions and antibodies. The eye lens is composed of long, thin “fiber cells” with elaborate plasma membrane ultrastructure stabilized by an actin/spectrin membrane skeleton (*45*). Palm2, in combination with Palm1 previously detected in fiber cells (*46, 47*), may be important for supporting this membrane skeleton, and it will be interesting to investigate whether the two isoforms segregate to distinct subcellular localizations in fiber cells.

All features of Palm2 immunoblotting and IF described above were confirmed with independent antibodies raised against non-overlapping immunogen sequences (rabbit anti-p21 and -p22, or mouse mAb 1A8), and their staining was markedly reduced or undetectable in tissues from *Palm2^em1(IMPC)J^* mutant mice processed in parallel. These mice were generated in the KOMP2 KO mouse project by a frameshift deletion, intending a null mutation. However, residual staining of strongly Palm2-immunofluorescent structures like the AIS, mossy fibers and sciatic NoR remained visible also in negative-control tissues from *Palm2^em1(IMPC)J^* mice (Fig. S1A1,B,C,F1). This led us to suspect that the KO of these mice might be leaky, and have residual expression of the C-terminal region of Palm2 against which the antibodies are directed. Indeed, immunoblot analysis of WT and mutant mouse brain homogenates at long development times indicated that while the 72 kDa Palm2 band was abolished, new weaker bands appeared in the mutant brain samples, detectable by two or even all three independent antibodies (Fig. S1H). The deletion of exon 3 in this mouse mutant, verified by us through PCR and sequencing of genomic DNA, eliminates the expression of full-length Palm2. However, it seems that aberrant splicing partially circumvents this deletion and the frame-shift it causes, and generates truncated Palm2 variants, one slightly longer and one shorter than intact Palm2. Retaining the C-terminal region of Palm2 which harbors the immunogen sequences recognized by the antibodies, as well as the membrane-anchoring CaaX motif, they are apparently still recruited e.g. to AIS, mossy fibers and sciatic NoR, where they give rise to residual specific staining in the tissues from mutant mice. Although not strictly a null mutant, we continue to refer to the *Palm2^em1(IMPC)J^* mice as “Palm2-KO” for brevity.

### Palm2 is enriched at the AIS of neurons in culture, complementary to the subcellular distribution of Palm1 along axons

In a previous study, we observed that Palm1 is a component of the MPS depleted from the AIS in hippocampal primary neurons (HPN) (*32*). Therefore, we turned to this experimental model to validate the histological data, and test whether Palm1 and Palm2 have complementary distributions along HPN axons. Line profiles of fluorescence intensity along the axon and dendrites confirmed that Palm2 was enriched in proximal axons, coinciding with the AIS marker ankG, whereas Palm1 was more abundant in middle and distal than in proximal axons (Fig. 2A-B2). Palm2 enrichment at the AIS was confirmed by all three antibodies (Fig. S2), and by CRISPR/Cas9-mediated knock-in of mEGFP at the N-terminus of Palm2 to visualize the endogenous Palm2 gene product (Fig. 2C). Both IF and knock-in approaches indicated that Palm2 was not restricted to the AIS but populated, at lower abundance, also distal axons and dendrites, consistent with the labeling of distal axons or neuropil in the hippocampal formation (Fig. 1B, Fig. S1A). Quantification revealed that Palm2 levels in the AIS were 2-3 times higher than in middle axons and dendrites (Fig. 2D). In comparison, ankG showed higher values for both compartments, indicating it was more stringently restricted to the AIS. In line with the histological data above, the anti-Palm2 immunoreactivity was reduced but not completely abolished in HPN from Palm2-KO mice (*Palm2^em1(IMPC)J^*; Fig. S3C).

**Figure 2:**
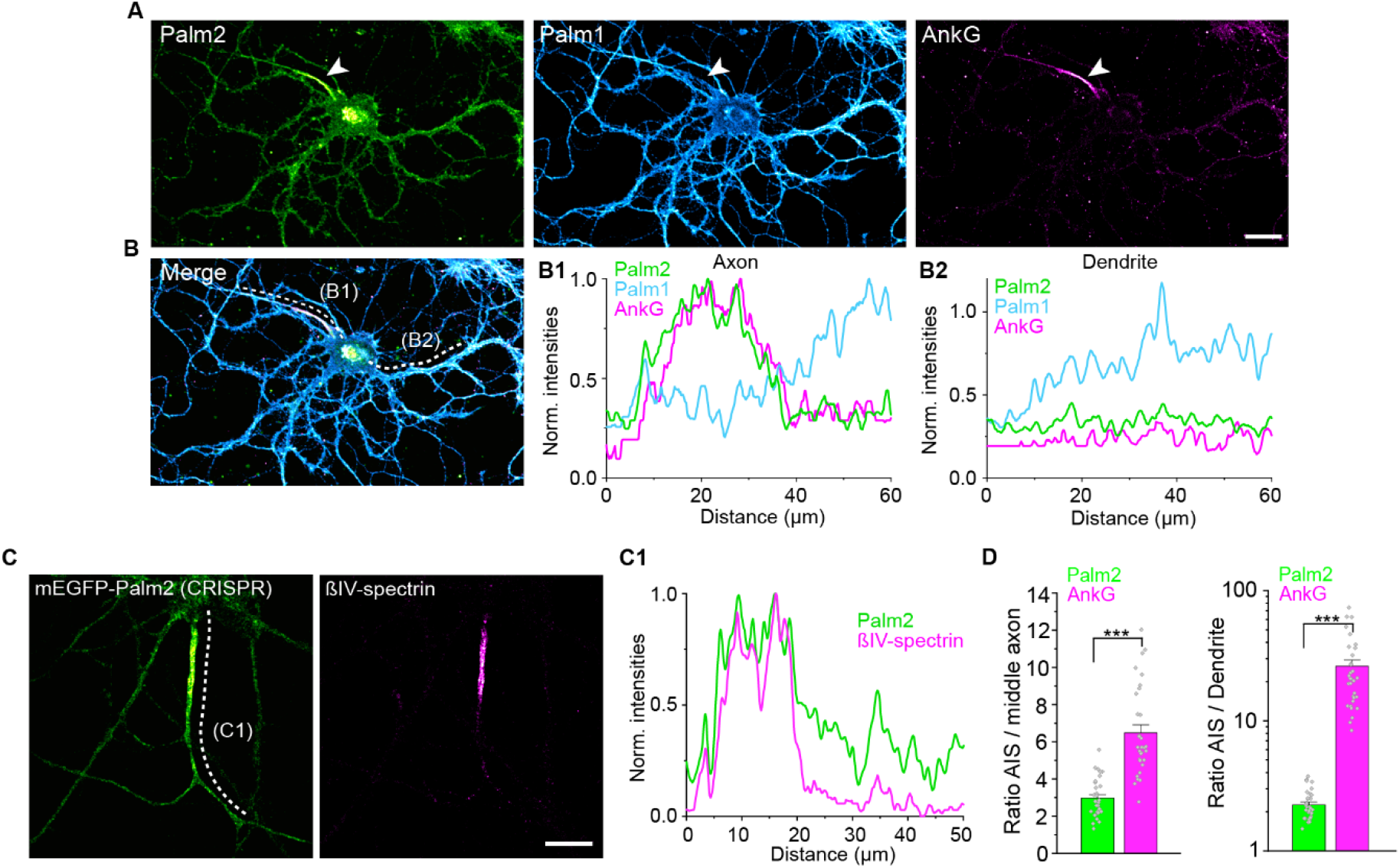
Palm2 is enriched in the AIS of primary hippocampal neurons. (A) Representative confocal images of rat HPN at DIV 19 immunolabeled against Palm2 (p21 antibody), Palm1 and ankG, and (B) merged image of all three channels. Images were processed using a Gaussian blur filter (σ=0.5 pixels). Scale bar: 20 µm. Dashed lines in (B) indicate smoothed and normalized fluorescence intensities of the proteins along an axon (B1) and dendrite (B2). Note that B1 and B2 are normalized to the same value, i.e. max intensity along B1 line profile. (C) Representative confocal image of a rat HPN at DIV 19 endogenously expressing mEGFP-Palm2 and detected with a nanobody against mEGFP (left), and corresponding βIV-spectrin AIS-labeling (right). Scale bar: 10 µm. (C1) Smoothed and normalized fluorescence intensities of mEGFP-Palm2 and βIV-spectrin along the axon indicated by the dashed line in (C). (D) Fluorescence intensity ratios of Palm2 (1A8) and ankG between the AIS/middle axon (left) and AIS/dendrites (right; note the logarithmic scale). (n_AIS/middle axon_ = 34 from N = 3 for both Palm2 and ankG, n_AIS/Dendrite_ = 33 from N = 3 for both Palm2 and ankG, rat HPN at DIV 19). All histograms show mean ± SEM. P-values in file S1.

Thus, we identified Palm2 as a new AIS component, consistent with recent AIS proteomic studies (*28–30*). Palm2 and Palm1 display complementary (though overlapping) subcellular distributions along the axon, similar to the complementary distributions of β-spectrin or ankyrin isoforms. These results from neurons in culture also demonstrate that the AIS enrichment of Palm2 is an intrinsic, cell-autonomous feature, independent of tissue context.

### Palm2 populates the AIS early in neuronal development, but displays no 190 nm periodicity

During neuronal development, the different AIS proteins appear at specific time points (*24, 30*). We therefore determined the subcellular localization of Palm2 during neuronal maturation in culture. Neurons at day in vitro (DIV) 1, 3, 5, 12 and 19 were co-stained against Palm2 and ankG, and the fluorescent signal obtained from the confocal images along the axons was quantified. At DIV 1, a weak Palm2 signal was present along the neurites and somas. From DIV 3 we observed Palm2 accumulation in the AIS, which remained constant from DIV 5 onward and colocalized with ankG (Fig. 3A,B). Given the relatively high intercellular variability of the signal, we analyzed if the abundances of ankG and Palm2 in individual axons correlate (Fig. 3C). This was not the case, *i.e.* high levels of ankG in an individual AISs did not correspond to high levels of Palm2, suggesting that the accumulation of the two proteins is mechanistically not closely linked. At all developmental stages, a weak Palm2 staining was also observed in the soma, dendrites and distal axons.

**Figure 3:**
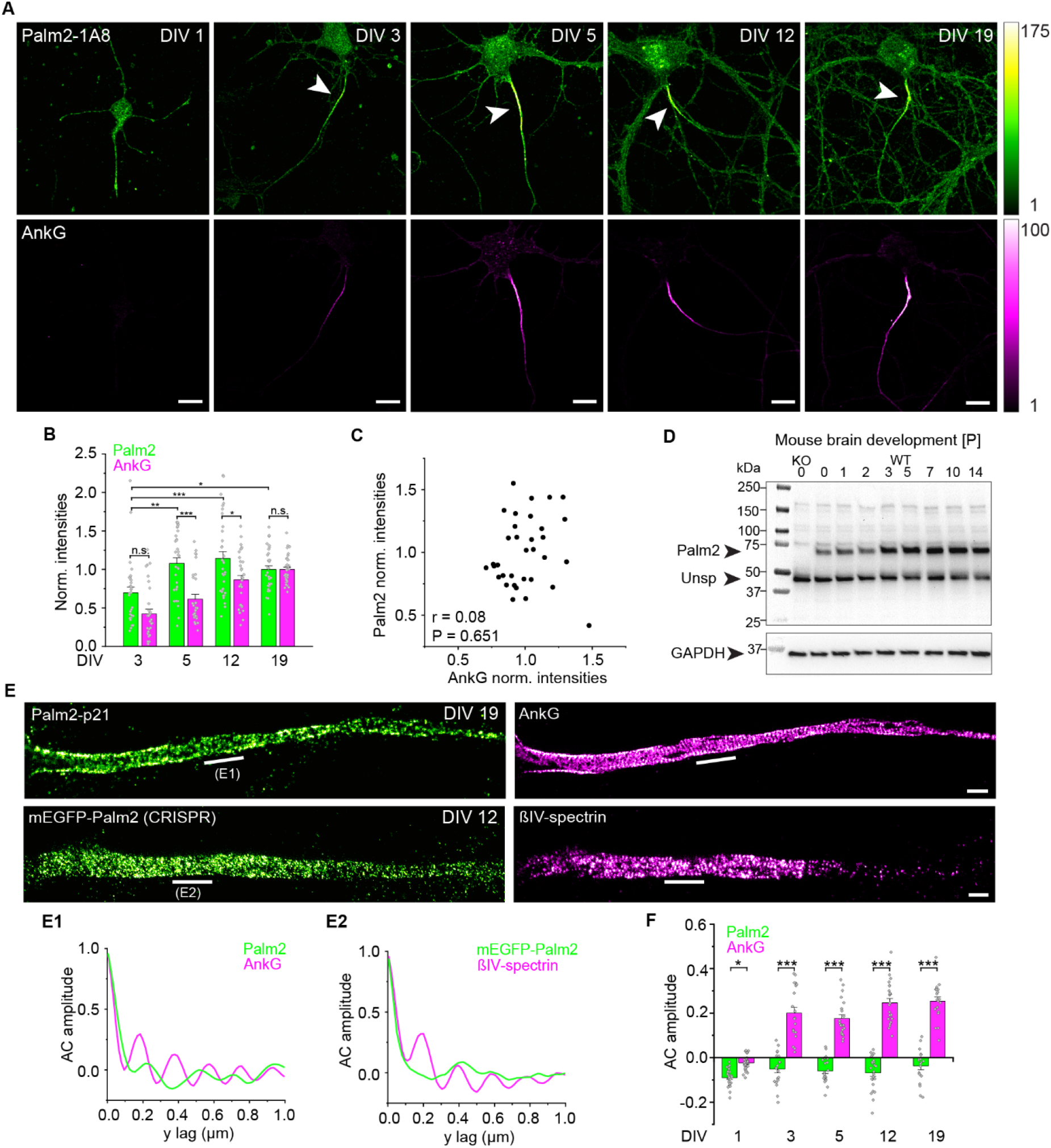
Palm2 is enriched at the AIS from early stages of neuronal development with a non-periodic nanoscale organization. (A) Representative confocal images of the distribution of Palm2 (1A8 antibody) and ankG in methanol-fixed rat HPN at DIV 1, 3, 5, 12 and 19. Fluorescence intensities are displayed for all panels as indicated by the color bars on the right. Arrowheads point at the AIS, identified by ankG labeling. Scale bars: 10 µm. (B) Normalized fluorescence intensities (A.U.) of both proteins within the AIS from DIV 3 to DIV 19. Axons analyzed from DIV 3 to DIV 19: 31 / 31 / 33 / 34 from N = 3. (C) Correlation scatter plot of the normalized fluorescence intensities of Palm2 versus ankG (DIV 19) shown in B. r, Pearsońs coefficient; P, P value. No correlation is observed. (D) Time course of Palm2 expression in postnatal mouse brain by immunoblot analysis. “Unsp”= unspecific band at ∼45 kDa typical for the 1A8 antibody, also detected in KO brain. Uncropped blot is shown in file S2. (E) Representative STED images of mature HPN double-immunolabeled against Palm2 (p21 antibody, DIV 19) (left; non-periodic) and the AIS-marker ankG (right; periodic), or against mEGFP-Palm2 knock-in (left; non-periodic) and βIV-spectrin (right; periodic, DIV 12). Lines in E indicate regions along which the AC analysis shown in E1 and E2 was performed. Scale bars: 1 µm. (F) AC amplitude analysis of endogenous Palm2 (p21 antibody) and ankG along the AIS over neuronal development. Axons analyzed for DIV 1 / 3 / 5 / 11-12 / 19-21: 28 / 20 / 22 / 24 / 19 from N = 3. Statistical analysis: One-way ANOVA with post hoc Tukey correction. All histograms show mean ± SEM. P-values in file S1.

We also analyzed the expression of Palm2 during brain development *in vivo*. A Western blot with an age series of samples from postnatal mouse brains (P0-P14; Fig.3D) showed that throughout this time span, only the standard Palm2-CaaX splice variant migrating at 72 kDa was detectable. Its signal was abolished in a sample from newborn Palm2-KO mouse, and its expression increased between P0-P7 and then leveled off. The 250 kDa variant (putatively Palm2-AKAP2) prominent in newborn mouse whole head (Fig. 1A) must therefore be expressed in the non-brain tissues, and more abundantly in neonatal head than in any of the adult tissues analyzed. We conclude that the properties of Palm2 characterized in this study, which were determined in HPN harvested from newborn rats or mice, are attributable to the 376-aa, lipid-anchored standard splice variant of Palm2 migrating at 72 kDa. The time course of Palm2 expression in whole brain is comparable to that observed for ankG and βIV-spectrin (*25*).

Next, we investigated the nanoscale organization of Palm2. All membrane-associated AIS markers characterized so far exhibit a periodic pattern in ∼190 nm intervals due to their association with the MPS (*1*). However, two-color STED nanoscopy revealed no periodic pattern for Palm2, independent of the fixation method (PFA, methanol or acetone), neuronal maturity, or the antibody used (Fig. 3E and Fig. S2B). In contrast, the AIS markers ankG, βIV-spectrin or Kv1.2, labeled in the same specimens by double-IF, displayed clear periodicities. These observations were analyzed using 2D autocorrelation (AC) and quantified by calculating the amplitude between the first peak at 190 nm and the average of the first 2 valleys at 95 nm and 285 nm (*24, 32*). As expected, ankG became periodically organized from DIV 3, whereas negative AC amplitudes were observed for Palm2 throughout neuronal maturation, indicating a lack of periodic nanoscale organization (Fig. 3F). STED nanoscopy of mEGFP-Palm2 knock-in neurons expressing the N-terminally tagged endogenous Palm2 also lacked detectable periodicity (Fig 3E, E2), ruling out the possibility that probing either the Palm2 N-terminus (mEGFP-tagged) or the C-terminal region (which is recognized by Palm2 antibodies) might be critical for the detection of periodicity.

In summary, Palm2 accumulates in the neurite that will develop into an axon even before we can detect ankG, and its AIS enrichment plateaus at an earlier developmental stage than ankG. However, at the nanoscale Palm2 is an untypical membrane-associated AIS component, in that no periodic pattern was observed regardless of neuronal maturity, fixation method, or labeling strategy.

### Palm2-KO neurons have shortened AIS and reduced excitability

Because Palm2 is present at the AIS from early stages of neuronal development, we wondered whether it is required for axon specification and AIS formation. For that, we investigated the development (DIV 1-15) of HPN from *Palm2^em1(IMPC)J^* mice. During early days *in vitro*, Palm2-KO neurons displayed a morphology similar to WT neurons, according to confocal microscopy and Sholl analysis. However, immature (DIV 3) Palm2-KO neurons displayed more processes longer than 120 µm (Fig. S3A,B). Interestingly, Palm1-KO neurons had shown the opposite phenotype, with fewer processes longer than 100 µm (see Fig. S5F of (*32*)). Later, Palm2-KO neurons developed normally in culture (Fig. 4A). Mature KO neurons (DIV 15) had formed axons of normal ankG staining lengths and intensities (Fig. 4B). With βIV-spectrin as marker, both AIS length (-12%) and intensity (-20%) were significantly reduced (Fig. 4C), but on the nanoscale no difference in AC amplitude of βIV-spectrin was observed between WT and KO neurons (Fig. 4C). These results indicate that Palm2 is not essential for axon specification, AIS formation or nanoscale organization, but its deficiency has gradual effects on βIV-spectrin accumulation and AIS length.

**Figure 4:**
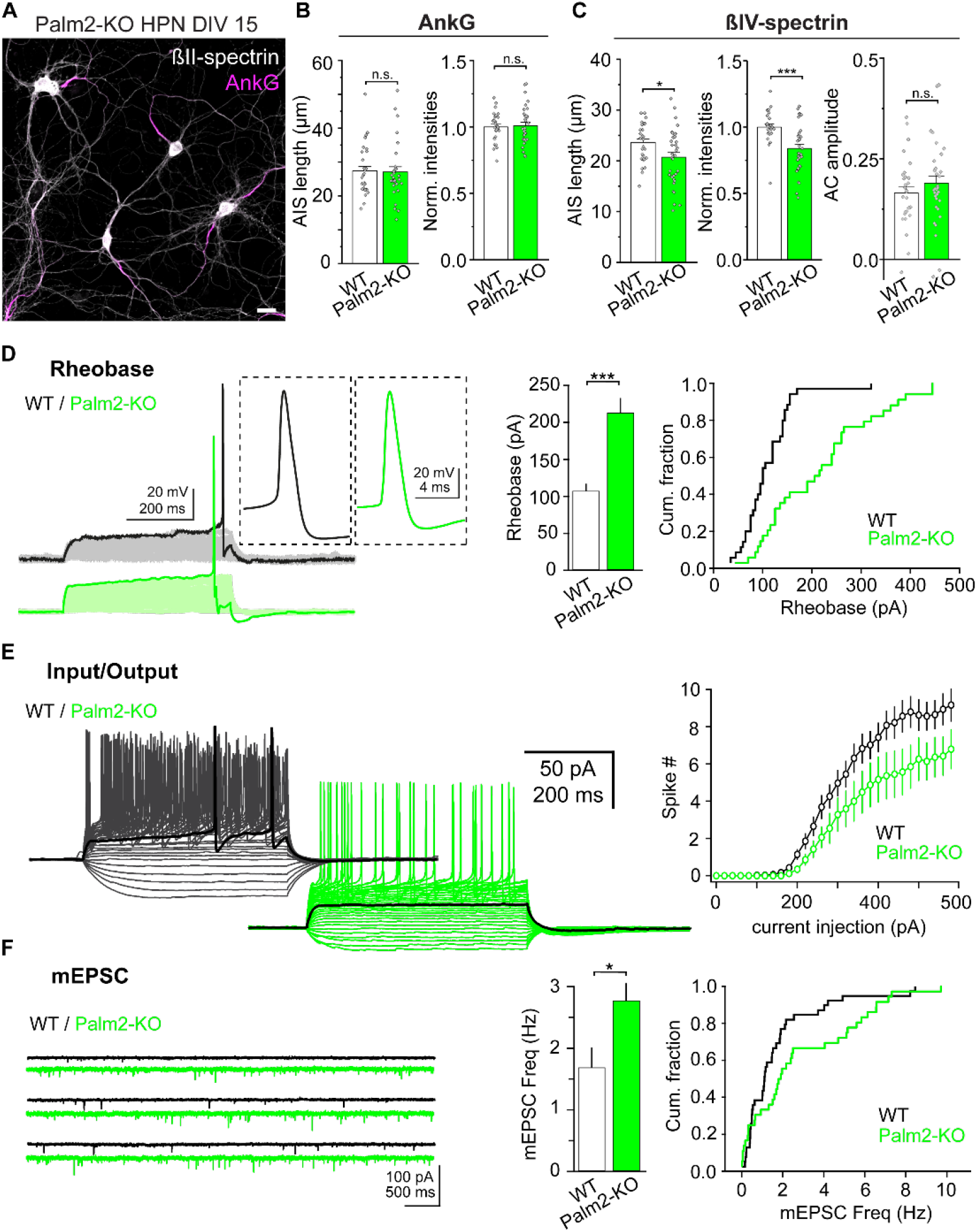
Palm2-KO neurons display a shorter AIS and altered electrophysiological properties. (A) Representative confocal image of mouse HPN derived from Palm2-KO at DIV 15. Scale bar: 20 µm. (B) AIS length (left) and local intensities (right) based on ankG staining in WT and Palm2-KO HPN at DIV 15. (C) Same as in B, but for βIV-spectrin, plus the AC amplitude analysis (right). (D) Representative rheobase measurements (5 pA steps for 500 ms) in WT and Palm2-KO neurons as well as the enlarged profile of an evoked spike. (Right) Averaged rheobase and cumulative distribution measured for both conditions. (E) Representative voltage responses of WT and Palm2-KO neurons upon current injections of increasing amplitude (25 pA steps for 500 ms from -100 pA to 400 pA). The responses to 200 pA current injection are highlighted in bold. (Right) Summary plot of the spike # as a function of current injected in both conditions. (F) Representative miniature EPSC (mEPSC) traces at -70 mV from WT and Palm2-KO mature neurons. (Right) Plots of the averaged mEPSC frequency (mean ± SEM) as well as the cumulative fraction in WT and Palm2-KO neurons. Statistical analysis for all histograms: Paired Sample T-Test. All histograms show mean ± SEM from n = 28-34, N = 3. P-values in file S1.

Next, we tested whether Palm2 depletion influences the electrophysiological properties of neurons (Fig. 4D-F, Fig. S3D-G). Although Palm2-KO neurons produced action potentials of similar height and half width as WT neurons, their overall excitability was strongly reduced. In particular, the rheobase (minimum current required to elicit an action potential) was significantly increased, while the number of spikes generated by an injected current (input/output curve) was lower in Palm2-KO neurons. Capacitance and input resistance were unaffected, arguing against gross changes in passive membrane properties or dendritic morphology. Furthermore, voltage clamp recordings revealed that Palm2-KO neurons displayed a higher frequency of miniature excitatory postsynaptic currents (mEPSC, Fig. 4F), whereas mEPSC amplitude and area were unaltered. Together, these data suggest that the reduced excitability of Palm2-KO neurons is likely a consequence of AIS shortening, rather than changes in dendritic morphology or passive membrane properties. The selective increase in mEPSC frequency but not amplitude, suggests a presynaptic increase in release probability or higher synapse numbers, potentially reflecting a homeostatic response to reduced intrinsic excitability.

Together, these results show that Palm2 is not essential for axon specification, AIS assembly, or action potential generation. However, its deficiency strongly affects several electrophysiological parameters. Strikingly, these effects of Palm2 deficiency are opposite to those induced by Palm1 depletion: Palm2 deficiency decreased axonal excitability but increased the synaptic mEPSC frequency, whereas Palm1 deficiency increased excitability while decreasing mEPSC frequency (*32*).

We showed above that the *Palm2^em1(IMPC)J^* mice, while unable to express full-length Palm2, do express a low background of truncated Palm2 derivatives, presumably generated by irregular splicing. The present findings might therefore be influenced by partial activities or dominant-negative effects of these aberrant splice products.

### Overexpressed Palm2 can integrate into the MPS

After characterizing Palm2-KO neurons, we went on to study the impact of Palm2 overexpression, electroporating wild-type HPN (DIV 0) with a plasmid encoding Palm2-CaaX N-terminally tagged with YFP. First, we investigated its impact on neuronal morphology, as well as on the subcellular and nanoscale organization of YFP-Palm2, at various stages of development. Overexpression approximately doubled Palm2 levels (Fig. 5A). Within one day, YFP-Palm2 populated distal neurites and filopodia where βII-spectrin was still absent (Fig. 5B-B2). As cells progressed to stage 3 (∼DIV 3, when the neurite that will become the axon outgrows the other neurites (*48*)), they displayed an increased branching of virtually all neurites, leading to a more complex neuronal morphology than in the control samples overexpressing cytosolic YFP (Fig. 5C,D). The peculiar morphology of immature neurons, with numerous processes and long filopodia, is an overexpression effect shared by all four membrane-anchored paralemmin isoforms, and is also observable in non-neuronal cell lines (*32, 49*). It probably reflects an interaction with actin but is likely unrelated to the MPS.

**Figure 5:**
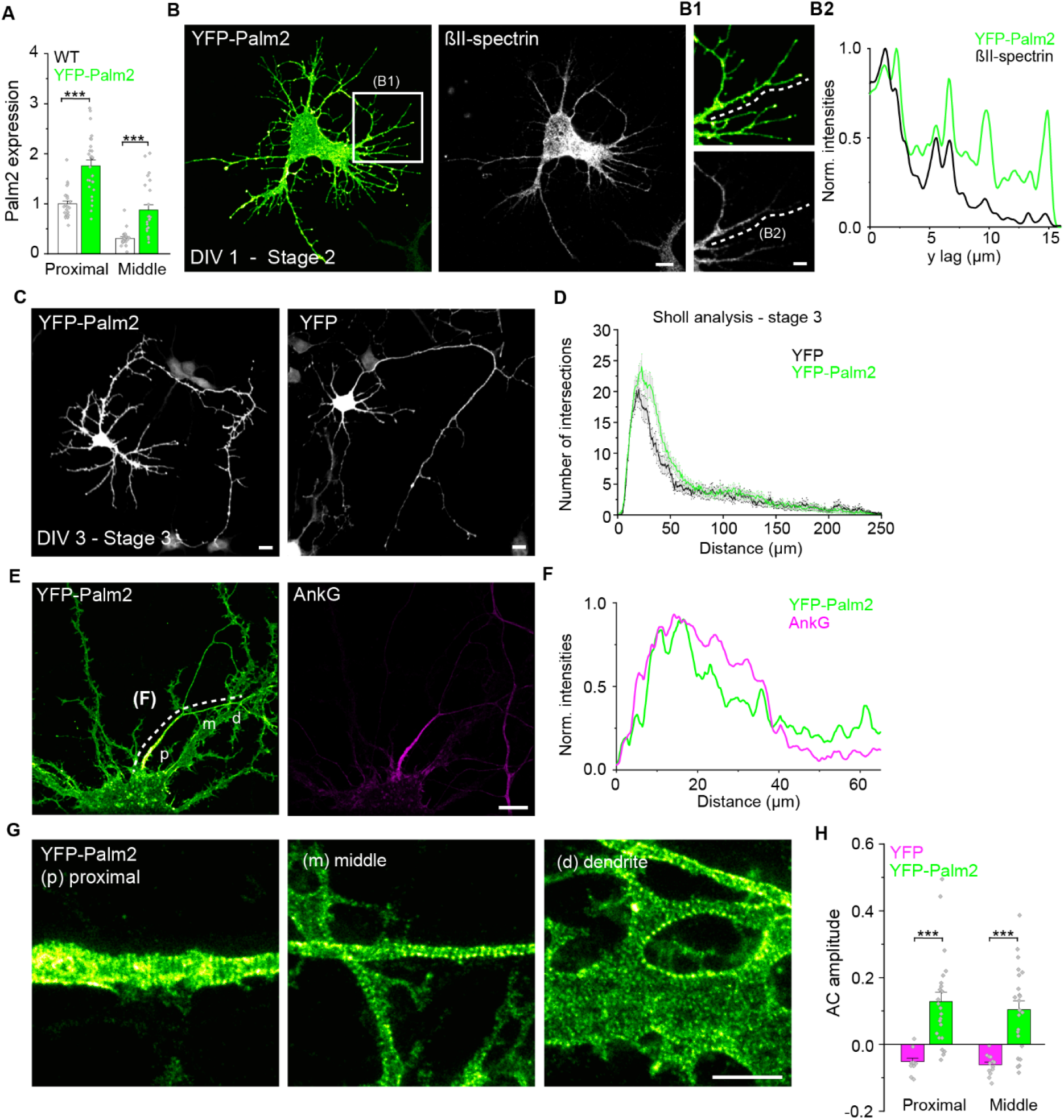
Overexpressed Palm2 alters neuronal morphology and assumes a periodic nanoscale organization. (A) Quantification of Palm2 overexpression after YFP-Palm2 electroporation. Axons analyzed in the proximal/middle regions: WT: 25/23; YFP-Palm2: 27/24. (B) Representative confocal images of a neuron overexpressing YFP-Palm2 at stage 2 of neuronal development, before axon specification. Scale bar: 10 µm. (B1) Close-up images of the region indicated by the white box show that Palm2 is already abundant at DIV 1 in distal neurites and filopodia, which are still poorly populated by βII-spectrin. Scale bar: 2 µm. (B2) Smoothed and normalized fluorescence intensities of Palm2 and βII-spectrin along the distal process indicated by the dashed line in the confocal image. (C) Representative confocal images of WT neurons at DIV 3 overexpressing either YFP-Palm2 (left) or YFP (right). Scale bar: 20 µm. (D) Sholl analysis of the neurons overexpressing YFP-Palm2 or YFP. Number of neurons analyzed YFP-Palm2 /YFP: 19/15, N = 3. (E) Representative confocal image of YFP-Palm2 and endogenous ankG in a rat HPN (DIV 19). A Gaussian filter of 0.5 was applied. Scale bar: 10 µm. (F) Smoothed and normalized fluorescence intensities of YFP-Palm2 and ankG along the axon as indicated by the dashed line in (E). (G) STED imaging of proximal (p), and middle (m) axonal, and dendritic (d) regions indicated in (E). Scale bars: 2 µm. (H) AC amplitude analysis of YFP and YFP-Palm2 in proximal and middle axons. Axons analyzed in the proximal/middle regions: YFP: 12/13; YFP-Palm2: 23/23. Statistical analysis: One-way ANOVA with post hoc Tukey correction. All histograms show mean ± SEM from N = 3. P-values in file S1.

In mature neurons (DIV 19), YFP-Palm2 was enriched at the AIS (Fig. 5E,F), demonstrating that recombinant Palm2, like endogenous Palm2, is recruited preferentially to this compartment, but also spreads further into the distal axon and dendrites. Analyzing the nanoscale organization of overexpressed YFP-Palm2 by STED, we were surprised to see a clear 190 nm periodicity in all neuronal compartments, including the AIS, middle and distal axons, and dendrites (Fig. 5G). AC amplitudes of YFP-Palm2 were similar in both AIS and middle axon (Fig. 5H), suggesting that it integrates in the MPS irrespective of the β-spectrin isoform. Of note, the absolute AC amplitude was still much lower than what was previously observed for Palm1 (∼0.1 for YFP-Palm2, vs ∼0.6 for YFP-Palm1, see Fig. S2G of (*32*)). This suggests that YFP-Palm2 is less efficiently expressed, or less efficiently recruited to the MPS than YFP-Palm1. Periodicity upon YFP-Palm2 overexpression was observed not only when detecting the N-terminal YFP-tag with an anti-GFP nanobody, but also when using the Palm2-1A8 antibody targeting the C-terminal region and thus labeling both recombinant and endogenous Palm2 (Fig. S4A). This demonstrates that the lack of periodicity of endogenous Palm2 is not due to Palm2 antibody performance, e.g. low affinity or less efficient access to C-terminal epitopes.

Next, we examined the localization of overexpressed YFP-Palm2 in relation to the MPS components βIV-and βII-spectrin, which are characteristic for the AIS, or the middle/distal axons and dendrites, respectively. We used antibodies that target the C-termini of both β-spectrins, half-way between the actin rings. YFP-Palm2 was found to be intercalated with the signals of both isoforms, as evident from cross-correlation analysis (Fig. 6A,B). For βII-spectrin, this intercalation was observed in AIS, middle/distal axons, and dendrites. This suggests that YFP-Palm2 assumes equivalent nanoscale positions, *i.e.* at or near the actin rings, in all subcellular compartments and relative to both β-spectrin isoforms. Together, these data indicate that higher levels of Palm2 through overexpression produce phenotypes similar to Palm1, including the ability to integrate into the MPS.

**Figure 6:**
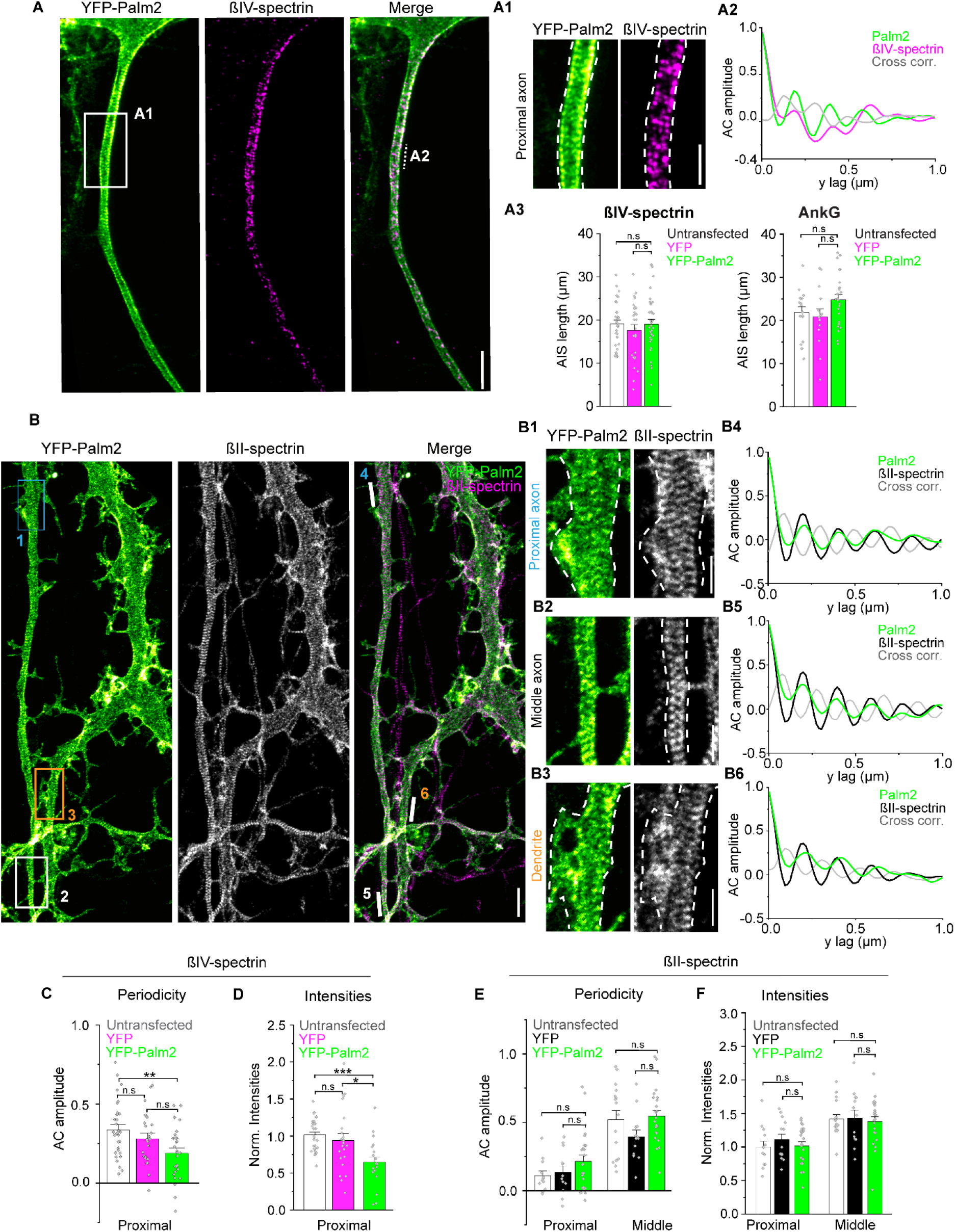
Palm2 overexpression impacts βIV-but not βII-spectrin periodicity and abundance. (A) Representative STED images of YFP-Palm2 and endogenous βIV-spectrin. Scale bar: 2 µm. (A1) Close-up of the proximal axon region indicated by the box in (A). Dashed lines indicate the axon outline based on YFP-Palm2 staining. Scale bar: 2 µm. (A2) AC and cross correlation amplitudes of YFP-Palm2 and βIV-spectrin calculated from the region indicated by the dashed line in the merged panel in (A). (A3) AIS length at DIV 19 based on βIV-spectrin (right) and ankG (right) stainings in untransfected neurons, or after overexpression of YFP or YFP-Palm2. Axons analyzed for βIV-spectrin/ankG: Untransfected: 32/17; YFP: 27/15; YFP-Palm2: 33/24 from N=3. Statistical analysis: One-way ANOVA with post hoc Tukey correction. P-values in file S1. (B) Same as (A) but for βII-instead of βIV-spectrin. A gaussian low pass filter of 1 pixel was applied to all STED images. Scale bar: 2 µm. (B1-B3) Close-up of proximal axon, middle axon, and dendrite regions indicated by the boxes in (B). Scale bars: 1 µm. (B4-B6) AC and cross correlation of YFP-Palm2 and βII-spectrin calculated from the regions indicated by the solid lines in the merged panel in (B). (C) AC amplitude analysis of endogenous βIV-spectrin along the AIS in untransfected neurons, or after overexpression of YFP or YFP-Palm2 at DIV 19. (D) Normalized fluorescence intensities (A.U.) of βIV-spectrin along the same axonal regions measured in (C). (E and F) same as C and D but for endogenous βII-spectrin. Axons analyzed in the C and D: Untransfected: 30; YFP: 23; YFP-Palm2: 27. In E proximal/middle: Untransfected: 12/16; YFP: 12/13; YFP-Palm2: 23/23. In F: Untransfected: 13/16; YFP: 13/13; YFP-Palm2: 23/23. Statistical analysis: One-way ANOVA with post hoc Tukey correction. All histograms show mean ± SEM from N = 3. P-values in file S1.

### Palm2 overexpression reduces βIV-spectrin periodicity and abundance, but does not affect βII-spectrin

Previously, we observed that overexpressed YFP-Palm1 not only integrated into the MPS, but even enhanced the periodicity of the endogenous MPS components βII-spectrin, adducin and ankB. Therefore, we wondered whether overexpressed YFP-Palm2 has a similar effect. AIS length, measured using both βIV-spectrin and ankG staining, remained unchanged following YFP-Palm2 overexpression (Fig. 6A3). However, although overexpressed YFP-Palm2 assumed a periodic organization in the AIS, it even reduced the periodicity of endogenous βIV-spectrin (indicated by reduced AC amplitude; Fig. 6C) and decreased its abundance compared to untransfected cells (Fig. 6D).

We then wondered whether overexpressed YFP-Palm2 has an effect on βII-spectrin, similar to Palm1 (*32*). However, both the nanoscale organization and the local abundances of endogenous βII-spectrin in proximal and middle axons remained unaltered upon overexpression of YFP-Palm2 (Fig. 6E,F). Therefore, while overexpressed YFP-Palm2 mimicked Palm1 to the extent of integrating into the MPS (though with a much lower AC amplitude), it failed to enhance the periodicity of both endogenous βIV-spectrin and βII-spectrin in middle and distal axons, which YFP-Palm1 achieves so strikingly (*32*).

### Palm2 binds β-spectrin/actinin isoforms with broad specificity

Our experiments demonstrated the ability of overexpressed Palm2 to interact with the MPS. We previously found that the “core domain” of Palm1 interacts with β-spectrin isoforms in the yeast-2-hybrid (Y2H) assay: to βII-spectrin with high avidity that resists up to 200 mM of the inhibitor 3-aminotriazol (3AT), but less strongly also to βI-, βIII-and βIV-spectrin (*32*). The core domain is conserved in Palm2 (aa ∼165-240, Fig. 7A,B), and we investigated therefore whether Palm2 also interacts with β-spectrin and related actinin isoforms. In Y2H tests, we matched mouse Palm2 (lacking exon 6; Fig. 7A, see Discussion) and human Palm2 (with exon 6) baits with preys encoding the Palm1-binding regions of β-spectrin isoforms I-V and the closely related actinin isoforms 1, 2 and 4 (Fig. 7C,D). Interactions were observed with βI-, βII-and βIV-spectrins which, however, were already suppressed by 2-5 mM 3AT (Fig. 7C). Human Palm2 gave slightly stronger interactions than mouse Palm2 and also interacted weakly with Actn1 and Actn4 (the non-muscle actinin isoforms; Fig. 7B). The presence or absence of exon 6 did not make a qualitative difference for the pattern of interactions, while the overall stronger interactions of hPalm2 may be due to higher expression of this bait in yeast (Fig. 7E). Also the corresponding Palm1-LexA bait fusion proteins were expressed at similar levels (Fig. 7E), so that the much stronger Palm1/βII-spectrin Y2H interaction is not due to higher Palm1 bait expression in yeast. In conclusion, mPalm2 and hPalm2 interact moderately with several β-spectrin/actinin isoforms, comparable in strength to the interactions of Palm1 with βI-, βIII-and βIV-spectrin (*32*). However, they lack the strong, preferential interaction of Palm1 with βII-spectrin that resists even 200 mM 3AT.

**Figure 7:**
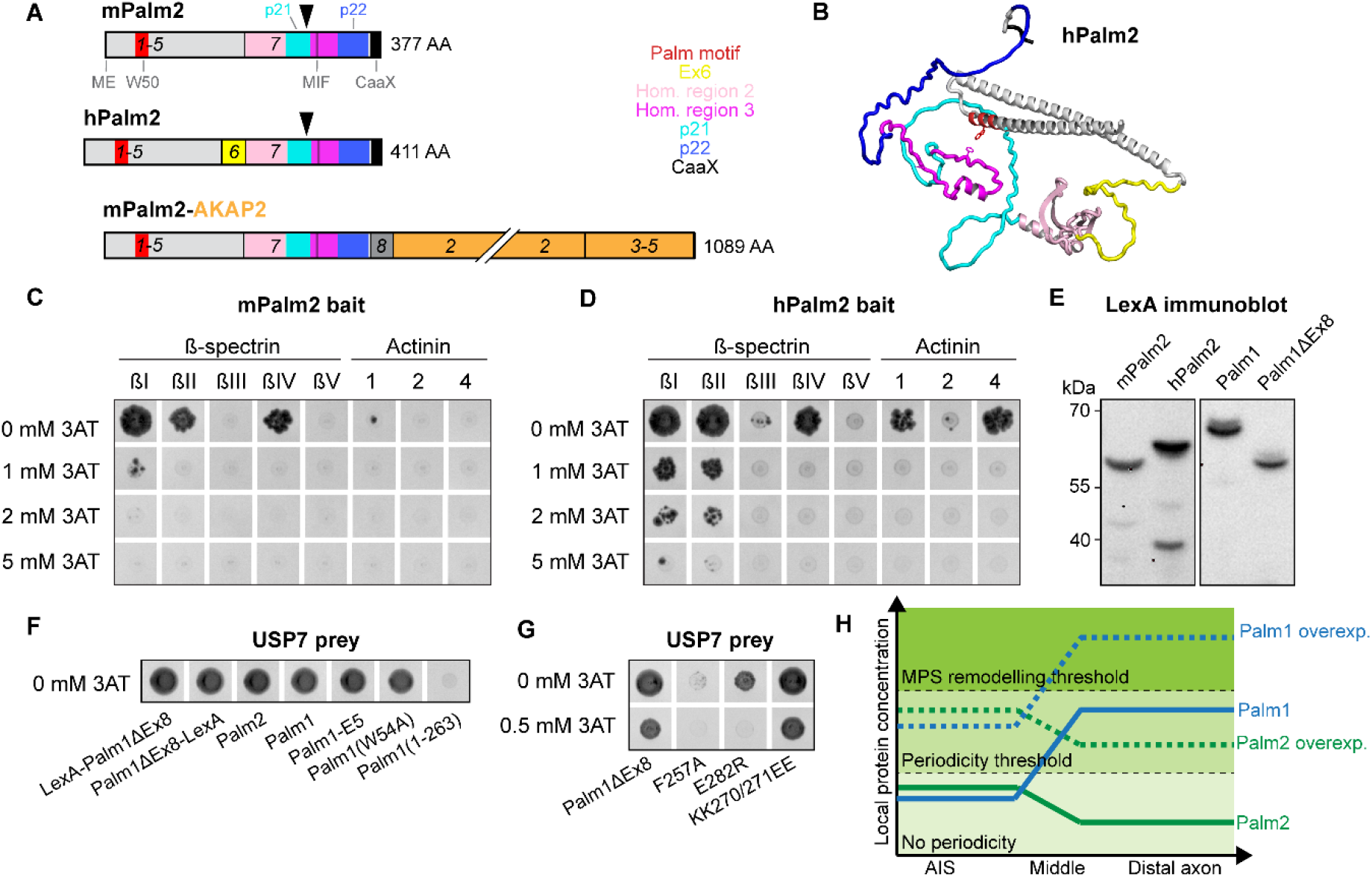
Pairwise Y2H analysis of paralemmin interactions. (A) Comparison of mouse and human Palm2, and of the chimera Palm2-AKAP2. Relevant regions are color-coded as in the caption on the right. Numbers indicate exon numbers. Black arrow shows where the baits used in C and D have been truncated to reduce autoactivation (see methods). Exon lengths not in scale with actual AA length. (B) Model of hPalm2 (Alphafold Q9Y2D5-8) with relevant regions highlighted as in A. Note that high likelihood of folding is predicted for the coiled-coil domain (homology region 1) and the core domain (homology region 2), whereas the other sequence regions are largely intrinsically unstructured (Movie S1). (C,D) Yeast spot colonies of mouse and human Palm2 baits show interactions with several members of the β-spectrin/actinin family. (E) Immunoblot analysis of the expression of Palm2 and Palm1 bait constructs in yeast, as used in experiments C-D and F-G, respectively. All lanes are from the same blot exposure, developed with anti-LexA-DBD. Full membrane shown in file S2. (F) The USP7 Ubl45 domain (RRSF…FEPQ) interacts as prey with various Palm1 and Palm2 bait constructs, but requires the C-terminal homology region 3 for this, deleted in Palm1(1–263). (G) Interaction of Palm1ΔEx8 with USP7 requires aa F257 and E282. All spot colonies are shown in the negative. (H) Model proposing different threshold levels at which Palm1 and Palm2 become periodic and are able to remodel the MPS (see discussion).

### Interaction of Palm2 and Palm1 with the de-ubiquitinase USP7 through the MIF motif suggests a link with proteostasis

Spectrins are unlikely to be the only paralemmin-binding proteins. A search for additional conserved binding partners was conducted using the Palm1ΔEx8 splice variant as bait, which does not bind spectrin in Y2H but shows strong periodicity when overexpressed (*32*). The screening of a human brain Y2H cDNA library produced several prey clones encoding the protease USP7. USP7 is a Ubiquitin-Specific Protease with many biological involvements including neuronal development, and its paralemmin-interacting C-terminal region, the tandem ubiquitin-like domains 4+5 (“Ubl45”), is critical for regulating its activity and specificity (*50, 51*). We verified this interaction by inverting the arrangement of the bait fusion protein: The library screen had been carried out with a bait fusion protein in which the LexA DNA-binding domain preceded the Palm1 sequence (LexA-bait) whereas the subsequent pairwise Y2H tests were performed in the inverse arrangement (bait-LexA) (Fig. 7F). Both arrangements of Palm1ΔEx8, and also Palm2, interacted with USP7 (Fig. 7F). We then explored the structural features of paralemmins responsible for binding USP7, employing deletion-and missense-mutant Palm1 bait constructs. The Y2H interaction was unaffected by the presence or absence of the exon-8-encoded spectrin-binding domain (full-length Palm1 vs. Palm1ΔEx8), the introduction of 10 phosphomimetic mutations (Palm1-E5), or the mutation of tryptophan 54 which abolishes the association of Palm1 with the MPS (Palm1(W54A)). Interaction was eliminated, however, by deleting the C-terminal third of Palm1 which encompasses “homology region 3” with the “MIF sequence motif” (Palm1(1–263)). We then introduced three missense mutations into Palm1ΔEx8, of amino acids which are conserved between Palm1, Palm2 and Palmd from multiple species: F257A, E282R and KK270/271EE (Fig. 7G). F257A abolished interaction with USP7 completely, E282R displayed weak residual interaction only at 0 mM 3AT, whereas the KK270/271EE mutant retained interaction up to 2 mM 3AT like the WT bait. Phenylalanine 257 is part of the “MIF” motif which is one of the sequence hallmarks invariably shared between Palm1, Palm2 and Palmd. This mutagenesis experiment was performed with Palm1 because its lack of Y2H autoactivation permits a better signal/background ratio (see Methods), but we expect its result to be applicable also to Palm2 and Palmd.

## Discussion

In the present study, we explored the role of Palm2 in the nervous system. Because of the scarcity of data on Palm2, we were guided by the known interactions of the other paralemmin isoforms with the actin-spectrin submembrane cytoskeleton. Palm2 is enriched at the AIS and NoR, with different localizations in NoR of the CNS (paranodal) and PNS (nodal). The founding member and most abundant isoform of the paralemmin protein family is Palm1, and we recently characterized Palm1 as a component and regulator of the MPS of neuronal axons (*32*). As Palm2 is the closest relative of Palm1, with 38% sequence identity and a collinear domain organisation, we expected it to be functionally analogous. The present results identify similarities but also differences between these paralemmin isoforms.

1. Palm2 and Palm1 are both attached to the inner face of plasma membranes through their C-terminal lipid anchors, and predominantly expressed in the nervous system. However, whereas Palm1 is virtually ubiquitous, Palm2 localization is more focused: it is enriched in the AIS and NoR, and in some non-myelinated axon populations like hippocampal mossy fibers.
2. Along axons, Palm2 and Palm1 display complementary distributions. Palm2 concentration is highest in the proximal axon, the AIS, whereas Palm1 concentration is higher in the more distal regions of axons. This intra-axonal distribution pattern parallels that of the βIV/βII-spectrin and the ankG/ankB ankyrin isoforms.
3. Endogenous Palm1 assumes the typical 190 nm periodicity of the MPS in middle/distal axons where its levels are highest, but it remains non-periodic in the AIS where its levels are lower. Also endogenous Palm2 is non-periodic in the AIS, despite being enriched there.
4. Overexpression of both YFP-Palm1 and YFP-Palm2 in immature neurons lead to a more complex neuronal morphology with more neurite branching and filopodial activity. Also in non-neuronal cells, Palm1 and Palm2 share similar properties, promoting cell migration and tumor metastasis (*52–54*). These more dynamic effects of paralemmin overexpression are probably MPS-unrelated, and may instead involve e.g. FERM proteins like ezrin (*52*).
5. On the nanoscale, overexpressed YFP-Palm1 and YFP-Palm2 both assumed a 190 nm periodicity in all neuronal processes including the AIS, indicating their capability to integrate into the MPS. However, whereas overexpression of YFP-Palm1 actively restructured the MPS and enhanced the periodicity of MPS-intrinsic proteins in the middle/distal axon (βII-spectrin, adducin and ankB), YFP-Palm2 did not have corresponding effects. On the contrary, YFP-Palm2 overexpression even reduced βIV-spectrin periodicity and abundance in the AIS, but did not perturb βII-spectrin in the AIS or more distally. This suggests that, in the context of the MPS, Palm2 preferentially interacts with βIV-spectrin and antagonizes it, even though in the Y2H assay it similarly binds to both spectrin isoforms.
6. Knock-out of Palm1 compromised the MPS nanoarchitecture without altering MPS-intrinsic protein levels. In contrast, the knock-out of Palm2 did not affect βIV-spectrin periodicity in the AIS, but led to lower βIV-spectrin abundance and shorter AIS.
7. Whereas Palm1 displayed an outstandingly avid interaction with βII-spectrin in the Y2H assay (and additionally weaker interactions with βI-, βIII-and βIV-spectrins), Palm2 lacked this strong preference for βII-spectrin and showed similarly moderate interactions with βI-, βII-and βIV-spectrin.
8. In spite of the complexity of the molecular and cellular effects described above, contrary electrophysiological phenotypes were very clear-cut: Palm2-deficient neurons displayed a decreased electrophysiological excitability (increased rheobase, reduced input/output ratio) and increased mEPSC frequency, whereas the Palm1-KO had the opposite electrophysiological effects.

Palm2 enrichment in the AIS, and its binding to β-spectrins in Y2H and to the MPS upon overexpression in cultured neurons, suggest that, analogous to the role of Palm1 in middle/distal axons, it may be important for modulating AIS assembly, homeostasis, and plasticity through its interaction with the submembrane skeleton. Manipulating Palm2 expression affects the AIS, but the finding that both Palm2 knock-out and overexpression reduce βIV-spectrin levels seems counterintuitive. An explanation may lie in the multi-domain architecture of paralemmins, with different domains affecting different aspects of the MPS (Fig. 7B, movie S1). The N-terminal coiled-coil domain (homology region 1) with the paralemmin motif around Palm2-W50 interacts with yet unidentified binding partner(s) located at the actin/adducin rings and, in Palm1, mediates its MPS-binding and remodeling function; the paralemmin core domain (homology region 2) interacts with spectrin; and homology region 3 around the MIF motif interacts with USP7, possibly affecting proteostasis. Different Palm2-concentration dependencies of these interactions may affect several AIS parameters differentially. For example, lack of Palm2 may lead to a less stable AIS scaffold, whereas enhanced Palm2 levels may cause higher USP7-mediated protein degradation. The coexistence of Palm2 with Palm1 at the AIS probably adds to the complexity of phenomena.

A delicate balance of protein levels could also underlie the lack of periodicity of endogenous Palm2, but also Palm1, in the AIS. As most membrane-associated AIS-enriched proteins are periodic, we were surprised that we could not detect this for endogenous Palm2 and invested much effort into verifying it. The simplest explanation may be that the local abundances in the AIS of both, Palm1 and Palm2, are below a critical threshold for associating detectably with the MPS, whereas the higher endogenous Palm1 concentration in the distal axon, or overexpression of either isoform, would lift the protein concentrations above this “periodicity threshold” also in the AIS (Fig. 7H). The absolute local concentrations of Palm2 or Palm1 in axons or AIS are unknown, but the Palm2 concentration in the whole brain is only 3% of Palm1. It is therefore possible that the absolute Palm2 concentration is still lower than that of Palm1, even in the AIS. Other factors such as local differences in molecular context (e.g. low βII-spectrin and adducin concentrations) or phosphorylation (both isoforms are multiply phosphorylated on up to 35 sites each) may contribute to the lack of periodicity of Palm1 and Palm2 in the AIS. Future research should investigate whether neuronal stimulation protocols that induce AIS remodeling also affect Palm2 periodicity or its phosphorylation. It is important to note that paralemmins could dynamically interact with the MPS even if they do not exhibit periodicity by STED. In immature Palm1-KO neurons, the lack of Palm1 was seen to impair βII-spectrin periodicity at a developmental stage when Palm1 itself was not yet detectably periodic in WT neurons (Fig. S5G of (*32*)). Even while still diffusely spread out over the plane of the plasma membrane, paralemmins could recruit MPS components and thus concentrate them near the plasma membrane. Moreover, a fraction of endogenous paralemmin molecules can already be associated with the MPS while periodicity is not yet detectable above the diffuse background. It is remarkable that a mere doubling of Palm2 concentration by overexpression has such a marked effect on its MPS association, suggesting a dynamic range that may extend into higher concentrations. In the future, it should be explored whether also endogenous Palm2 displays detectable periodicity in histological structures where it is particularly abundant, e.g. in hippocampal mossy fibers, the indusium griseum, or the nodal gaps of sciatic NoR. It also remains to be investigated whether yet higher expression levels of recombinant Palm2, exceeding those that we achieved in the present experiments, can pass a “remodeling threshold” (Fig. 7H) and enhance the periodicity of endogenous MPS proteins, or whether Palm2 is intrinsically lacking this propensity of Palm1.

It is noteworthy that during development, Palm2 is detectable in the axon even before ankG, and it is enriched in the AIS from DIV 3, as early as ankG, the accepted master organizer of the AIS. This is reminiscent of the early colonization of distal axons by Palm1, preceding the establishment of the axonal MPS. However, as Palm2-KO neurons still develop an ankG-positive AIS, the recruitment mechanisms of the two proteins do not seem to be closely linked. The AIS and NoR share many molecular constituents, and we demonstrate here that Palm2 is among them. Uniquely, however, Palm2 co-localizes with ankG at the nodal gaps in only a subset of axons of sciatic nerves, apparently of sensory neurons, but not in the CNS, where it is instead present at the paranodes. While the detection of Palm2 at only a subset of NoR nodal gaps in the PNS could be attributed to differential Palm2 expression levels in sensory and motor neurons, the absence of Palm2 at ankG-positive nodal gaps in the CNS demonstrates that Palm2 is not an obligatory companion of ankG. As Palm2 mRNA is highly expressed in oligodendrocytes (https://brainrnaseq.org/), paranodal Palm2 might localize to the myelin loops rather than to the axon. Interestingly, these loops contain ankG in the CNS but ankB in the PNS (*11*). Determining the exact localizations of Palm2 in CNS-and PNS-NoR (axonal or glial, periodic or not) is required to develop a cell-biological understanding of the targeting and the functions of Palm2 at NoR.

The molecular features and interaction partners of Palm2 involved in its targeting to the specific subcellular compartments, remain to be determined. The Palm2-specific sequences which flank homology region 3 and which we chose as immunogens to generate Palm2-specific antibodies (p21 and p22), are prime candidates for this (Fig. 7B, note that the immunogen used to generate 1A8 contains the p22 sequence). The deletion of exon 3 in the *Palm2^em1(IMPC)J^* mice removed the paralemmin motif around W50, but residual truncated expression products are still targeted to AIS and sciatic nodal gaps, suggesting that C-terminal Palm2 sequences are responsible for this targeting. Particularly the p22 sequence immediately adjacent to the membrane-anchored C-terminus (aa 314-366; KSLR…PGTQK) is an attractive candidate for mediating the recruitment of Palm2, and also its aberrant splice products in the *Palm2^em1(IMPC)J^* mice to the AIS and NoR, by binding to neighboring membrane proteins. Whereas the p21 sequence displays moderate inter-species sequence conservation (82/45/41% aa identity between mouse and human/chicken [Gallus gallus]/frog [Xenopus laevis]), p22 sequence conservation is exceptionally high (98/89/83% identity between mouse vs. human/chicken/frog). This indicates strong selective pressure on the p22 sequence, whereas e.g. the collinear C-terminal sequence in Palm1 is virtually non-conserved between mouse vs. chicken/frog and presumably has only spacer function. If this C-terminal spacer sequence of 50-60 aa elevates Palm1 above the layer of intrinsic membrane proteins whereas the p22 sequence of Palm2 is embedded between them, Palm2 may vertically stitch the actin-spectrin submembrane skeleton more tightly to the AIS membrane than Palm1. Horizontally, sandwiched between its protein neighbors, it may contribute to the diffusion barrier function of the AIS.

The different degrees of inter-species sequence conservation also imply that the anti-p22 antibody is likely to detect Palm2 in most or all vertebrates, whereas anti-Palm2-p21 cross-reacts with guinea-pig (Fig. S1G) and probably other mammalian but perhaps not with non-mammalian Palm2 orthologs. The p22 sequence is rich in serine/threonine (26%) as well as aspartate/glutamate residues (19%), all conserved between mouse, chicken and frog. Eight closely spaced serine/threonine residues in p22 are phosphorylated (S315, S341, T343, T344, S347, S348, S355, S358; https://www.phosphosite.org/proteinAction.action?id=31726300&showAllSites=true), reminiscent of the ankyrin-binding sequences of Nav1 sodium channels and of the scaffold protein IQCJ-SCHIP-1, which are also D/E-as well as S/T-rich and mediate the targeting of these proteins to the AIS, dependent on phosphorylation by the AIS-enriched protein kinase CK2 (*55, 56*). As the prenyl-dipalmitoyl membrane anchors of paralemmins preferentially associate with lipid raft-like microdomains (*57*), local membrane lipid composition may also contribute to the targeting.

Palm2 immunodetection in some tissue structures (AIS, mossy fibers) depended upon the fixation/permeabilization conditions (PFA/Triton/heat-induced antigen retrieval vs. fresh-frozen/acetone), and also differed in sensitivity to the antibodies (p21, p22, 1A8). Most strikingly, AIS staining in tissue sections was strong with fresh-frozen/acetone-postfixed but not with PFA/Triton/heat-treated specimens, whereas mossy fibers and other hippocampal axon populations were stained only in PFA/Triton/heat-treated specimens. NoR in the CNS, or lens fiber cells, stained similarly under both conditions of specimen processing. We suppose that the different epitopes are differently masked by the molecular context in the respective subcellular localizations, and differentially exposed by different conditions of fixation and permeabilization. All immunogen sequences lie close to the C-terminus of Palm2, and close to the plasma membrane due to the C-terminal lipid anchor, and they might be easily covered by their interaction partners and other membrane proteins. Therefore, the localization pattern of Palm2 in tissues may reflect actual differences in local Palm2 concentrations, but also epitope accessibility depending on molecular context, or epitope phosphorylation at the densely clustered serine/threonine residues in the p22 sequence. In the present discussion, we imply for the sake of brevity that absent Palm2 immunostaining indicates the absence of the protein. However, the observation that Palm2 detection with the existing antibodies can be highly sensitive to the technical conditions, cautions that lack of immunostaining may also be due to, e.g., local molecular context or epitope phosphorylation. Also Zhang et al. (*10*) discuss that immunodetection of Kv1 and K2P potassium channels at the AIS critically depends on fixation conditions.

Palm2 occurs in several splice variants and differential promoter transcripts, which must be considered for the interpretation of the present results (Fig. 7A). (1) The standard variant is collinear in aa sequence with Palm1, featuring a C-terminal CaaX motif which becomes modified with a prenyl-dipalmitoyl membrane anchor (*49, 52*). This is the only detectable variant in adult and postnatal brain of mice, which therefore underlies the properties of Palm2 that we describe in the present study in rodent brain and cultivated neurons, and which we employed as YFP-Palm2 in the overexpression experiments. (2) Natural fusion proteins of Palm2 sequences with a little-studied protein kinase A-binding protein encoded by a downstream gene (AKAP2, a.k.a. PAKAP), that replaces the CaaX membrane anchor, can be detected in several tissues (higher in neonatal than in adult ones; Fig. 1A), but do not affect the present results as they were undetectable in the brain. (3) A subgroup of rodent species (including mice, rats and hamsters) is genetically peculiar in that they express Palm2 without exon 6 (exon numbering of (*44*); termed exon F2 in (*31*)), resulting in a mouse Palm2 protein of 376 aa (Fig. 7A). These species have lost Palm2 exon 6 from their genomes, whereas most other rodents and all other mammals (including guinea-pigs and humans) have retained it (human Palm2 is 411 aa long). Therefore, the characterization of endogenous mouse and rat Palm2 and recombinant mouse YFP-Palm2 in the present study involved the variant lacking exon 6. However, in several experimental paradigms we also analysed the guinea-pig or human Palm2 variants with exon 6 in comparison, but observed no marked differences to the mouse variant: expression levels of mouse and human bait proteins in yeast (Fig. 7E); Y2H interaction with β-spectrin/actinin isoforms (Fig. 7C-D); AIS localization in mouse and guinea-pig brain (Fig. S1G). (4) Palm2 expression seems to involve two previously unnoticed alternative transcription starts. The originally characterized Palm2 begins with the aa sequence MAEAEL… encoded by exon 1 (“exon B” in (*31*)), and the aa numbering in the present study also refers to this amino terminus. However, one of three murine expressed sequence tags (ESTs) and 6 of 7 human ESTs in the sequence database, encode an N-terminus with two additional aa (<u>ME</u>MAEAEL…) preceded by a different 5’-untranslated sequence, which are encoded by a previously unrecognized “exon zero” (a.k.a. “exon A”) located 135 kb upstream of exon 1 in the mouse genome. Of note, Palm1 and Palm3 also possess exon A counterparts (*31*), such that the identification of a Palm2 exon A underpins the high degree of homology between the paralemmin isoform genes. We suspect that the presence or absence of the additional two N-terminal aa in Palm2 has little or no impact on the function of the protein (they are not included in YFP-Palm2), but that the use of promoters A or B is probably important for expression control of Palm2 and its downstream splice variants like Palm2-AKAP2. However, we cannot strictly exclude that the N-terminal tags introduced in our experiments ([ME]MAEAEL… in endogenous Palm2, to YFP-AEAEL… in the expression plasmid or [ME]M-mEGFP-AEAEL… in the CRISPR/Cas9-KI) might affect functional aspects of the N-terminal region, though the association of YFP-Palm2 with the MPS is not obstructed. Important for the present study, the CRISPR/Cas9-KI of the rat Palm2 gene placed the mEGFP tag in exon B, so that the expression products are predicted to carry the mEGFP tag, regardless of whether they are expressed from promoter A or B.

The N-terminal coiled-coil domain, and the paralemmin motif around Palm2-W50 embedded in it (Fig. 7A,B), is conserved between all four paralemmin isoforms, and mutation of Palm1-W54 abolished the ability of Palm1 to bind and remodel the MPS (*32*). Identifying its mode of binding at or near the actin rings will provide decisive insight for understanding how paralemmins interact with the actin/spectrin cytoskeleton. However, in several Y2H screens with all paralemmin isoforms under a variety of conditions, we were unable to identify ligands for the coiled-coil domain. Perhaps it binds to a composite interface to which several proteins of the actin rings and/or of the adjacent parts of α/β-spectrin contribute, making it undetectable by Y2H library screening which presents only individual prey polypeptide sequences of limited length to the bait. Binding of the coiled-coil domain to a composite interface would directly explain how Palm1 enhances the organization of the MPS by connecting several of its proteins, and other approaches like co-immunoprecipitation, chemical crosslinking or proximity labeling may be needed to identify the contributing proteins. Adducin and protein 4.1G, which co-immunoprecipitated with the paralemmin isoform, Palmd, may be among them (*35*).

Our detection of Y2H interactions of Palm1 and Palm2 with the de-ubiquitinase, USP7, suggests that, by recruiting USP7 to the AIS, the NoR and the MPS, paralemmins could integrate mechanisms of proteostasis into these compartments. The sequence overlap of the Palm1-interacting human USP7 clones detected in our Y2H screen, aa 891-1066 (RRSF…FEPQ), closely matches the tandem ubiquitin-like domains 4 & 5 (“Ubl45”) known to be crucial for USP7 activity and specificity (*50, 51*). Recent proteomic studies also detected USP7 at the AIS (*28, 30*). Another proteomic study identified a link between Palm2 and another putative de-ubiquitinase, OTUD7A, which in turn also interacted with βII-and αII-spectrin as well as ankB and ankG (*58*). Lastly, the proteasome adapter Ecm29 was identified at the AIS (*59*), and both NMDA-as well as neuronal injury-induced AIS plasticity require the ubiquitin-proteasome system (*27, 60*). In the present work, we did not pursue the potential link between paralemmins, USP7 and the proteostasis of the submembrane skeleton further, but this deserves future research attention and is likely relevant in the emerging field of AIS plasticity. The missense mutations in the MIF motif of homology region 3 which abolish the paralemmin-USP7 interaction, strongly support the specificity of this interaction and will enable cell-biological experiments to pursue this further.

We conclude that Palm2 is a novel constituent of the AIS and NoR, with unique characteristics in terms of a “conditional” association with the MPS and a localization to different compartments of central (paranodal) vs. peripheral (nodal) NoR. Beyond the AIS and NoR, Palm2 is enriched in select axon populations like hippocampal mossy fibers, and in the fiber cells of the eye lens. Genetic Palm2 deficiency reduces AIS length and axonal excitability. At the molecular level, Palm2 binds several β-spectrin isoforms and the de-ubiquitinase USP7. These results indicate that Palm2 interacts with the submembrane actin-spectrin cytoskeleton, similar to its isoforms Palm1 and Palm3, and we expect that its further characterization will advance our understanding of the assembly, remodeling and functioning of the AIS and the NoR.

## Materials and methods

### Palm2 antibodies

Isoform-specific Palm2 antisera were raised in rabbits at Pineda Abservice (Berlin, Germany) against the non-overlapping mouse Palm2 sequences Palm2-p21 (aa 207-273; GQSS…NLDQ) and Palm2-p22 (aa 314-366; KSLR…PGTQK), GST-tagged in pGEX-4T-2 (SmaI site), expressed in bacteria and purified by glutathione affinity chromatography. These immunogen sequences were chosen for their lack of sequence similarity with the other paralemmin isoforms, flanking homology region 3 in the C-terminal half of Palm2 (Fig. 7A-B, movie S1). Sera were affinity-purified with the respective, immobilized immunogen fusion proteins. Mouse monoclonal antibody M09 (clone 1A8), raised against the human Palm2 immunogen sequence aa 321-411 (EDEEE…CCVVM), was purchased from Abnova (Taiwan, cat. H00114299-M09).

### Palm2-mutant mice

Mutant mice were generated at the Jackson Laboratory (JAX) as part of the KOMP2 project (strain name: C57BL/6NJ-*Palm2^em1(IMPC)J^*/Mmjax or C57BL/6NJ-*Pakap^em1(IMPC)J^*/Mmjax; stock number: 042398-JAX). Mutagenesis had been performed by injecting Cas9 RNA and 4 guide sequences CATGGCTTTCGGTATCTGCT, AGACAACAGATGACACCTTT, CGTGCTGCGACTTTTCCCTG and TCCTAAATAGATACTGGTGT, which resulted in a 292 bp deletion beginning at Chromosome 4 negative strand position 57,648,243 bp and ending after 57,647,952 bp (GRCm38/mm10). This mutation deletes ENSMUSE00001288752 (exon 3) and 161 bp of flanking intronic sequence including the splice acceptor and donor and is predicted to cause a change of amino acid sequence after residue 42 and early truncation 10 amino acids later through a reading-frame shift. The genotyping protocol is available from the MMRRC web-site: https://www.mmrrc.org/catalog/sds.php?mmrrc_id=42398. Mice were reconstituted from frozen sperm obtained from JAX, at the Transgenic Facility of the Max Planck Institute for Multidisciplinary Sciences by *in-vitro* fertilization of C57BL/6N mouse eggs. The genome sequence around the exon 3 deletion was PCR-amplified by us, and the mutation verified by sequencing. Homozygous Palm2-mutant mice were healthy and fertile with no obvious phenotype. Mice were bred and kept in the mouse facility of the Max Planck Institute for Multidisciplinary Sciences (City Campus). All regulations given in §4 Animal Welfare Law of the Federal Republic of Germany (§4 TierSchG, Tierschutzgesetz der Bundesrepublik Deutschland) were followed. No specific authorization or notification was required for the breeding of animals, or for the procedures performed in this study. We have complied with all relevant ethical regulations for animal use.

### IF of mouse and guinea pig brain and eye tissues

Animals were sacrified by decapitation under isoflurane anaesthesia, brains, cerebella and eyes excised within minutes, and either immersion-fixed for 24 h in 4 % paraformaldehyde (PFA) in phosphate-buffered saline (PBS) at 4°C, or fresh-frozen in TissueTek on dry-ice and stored at -80°C until cutting. PFA-fixed eyes were covered with TissueTek and frozen at -80°C until cutting. PFA-fixed eyes and fresh-frozen brains and cerebella were cut into 12 µm thick sections using a cryostat (Leica), mounted on superfrost plus slides and stored at -80°C until staining. PFA-fixed brains and cerebella were cut into 50 µm thick sections using a vibratome (Leica) and stored in a cryoprotective solution (25 % glycerol, 25 % ethylene glycol, 50 % PBS pH 7.4) at -20°C until staining. PFA-fixed brain and cerebellum sections were stained free-floating, PFA-fixed eye and fresh-frozen brain sections were stained on slides. All were washed in Tris-buffered saline (TBS; 50 mM Tris, 150 mM NaCl, pH 7.2) and incubated in antigen-retrieval citrate buffer (10 mM citrate, pH 6.0) overnight at 60°C. The sections and slides were then washed in TBS and blocked for 1 h in 10 % normal goat serum and 0.3 % Triton X-100 in TBS at room temperature (RT). Primary antibodies were diluted in 5 % normal goat serum and 0.3 % Triton X-100 in TBS. Sections and slides were incubated overnight at 4°C, followed by washing and incubation with the respective secondary antibody for 1 h at RT, and counterstaining with DAPI. After staining, the sections were washed in TBS, dried and mounted with Entellan (Sigma-Aldrich). Fresh-frozen sections were post-fixed with acetone for 10 min at -20°C. After drying, they were washed with TBS and blocked for 1 h in 10 % normal goat serum and 0.3 % Triton X-100 in TBS at RT. Primary antibodies were diluted in 5 % normal goat serum and 0.3 % Triton X-100 in TBS and sections incubated overnight at 4°C, followed by washing and incubation with the respective secondary antibody for 1 h at RT, and counterstaining with DAPI. After staining, the sections were washed and covered with AquaPoly Mount (Polysciences). Primary marker antibodies were: mouse anti-ankyrin G (Synaptic Systems, cat. 386 011, 1:500), chicken anti-ankyrin G (Synaptic Systems, cat. 386 006, 1:500-1:1000). Secondary antibodies were: donkey anti-rabbit Cy3 (Jackson ImmunoResearch, cat. 711-165-152), goat anti-rabbit Cy3 (Jackson ImmunoResearch, cat. 111-165-144), donkey anti mouse Alexa Fluor 647 (Jackson ImmunoResearch, cat. 715-605-151), donkey anti-chicken Cy5 (Jackson ImmunoResearch, cat. 703-175-155).

Imaging of mouse and guinea pig tissues (Fig. 1 and S1, except 1D,E,F, and S1D,E) was performed using an Axio Observer 7 microscope (Zeiss) equipped with an apotome 3, Zeiss Axiocam 807 mono, X-Cite XYLIS lamp (Excelitas) and the following objectives: Fluar 2.5x/0.12, Plan-Apochromat 10x/0.45, Plan-Apochromat 20x/0.8, Plan-Apochromat 63x/1.4 Oil. The images were processed using the Zen software 3.6.

For data shown in Fig. 1D and Fig. S1D+E (brain NoR), mice were transcardially perfused with 4 % PFA in PBS. After dissection, brains were post-fixed for 4 h in the same 4 % PFA solution, and then transferred to 30 % sucrose (in PBS) overnight. Brains were cut into ∼30 µm thick sections using a cryo-microtome. Free floating sections were washed 3 times in PBS before antigen retrieval in sodium citrate buffer (10 mM sodium citrate, 0.05 % Tween20, pH 6) at 80°C for 20 min, followed by three further washes 10 min each in PBS. Samples where then incubated for 1-2 hours at RT in PBS supplemented with 0.3 % Triton X-100 and 3 % BSA at RT. Primary antibody incubation was performed in the same buffer ON at 4°C in constant agitation (Palm2 1A8, 1:1000; anti-ankyrin G Synaptic Systems, cat. 386 003, 1:1000). Following three washes in PBS, incubation with secondary antibodies was performed for 1-2 h at RT in PBS with 0.1 % Triton X-100 (anti-mouse STAR580, Abberior ST580-1001-500ug, 1:500; anti-rabbit Alexa Fluor 488, Thermo Fisher, cat. A11034, 1:500). Lastly, sections were washed and deposited on coverslips, air-dried, and mounted in Mowiol supplemented with DABCO. Confocal imaging was performed on an Abberior Expert line microscope described below.

### IF of sciatic nerves

C57BL/6N mice (female, 92 weeks) were sacrificed by decapitation under isoflurane anaesthesia, their sciatic nerves immediately extracted, and fixed in 4 % PFA in PBS on ice for 50 min. To quench free aldehyde groups, the nerves were then incubated in PBS supplemented with 100 mM glycine and 100 mM NH_4_Cl for additional 50 min, and stored in PBS at 4°C. After partial teasing, free floating fibers were permeabilized by incubation in methanol on ice for 30 min, followed by blocking with 1 % BSA in PBS for 45 min at RT. Primary antibody incubation was performed using a combination of anti-Palm2-p22 (1:200) and anti-CASPR (NeuroMab, cat. 75-001, 1:200) in PBS with 0.05% Triton X-100. The primary antibodies were incubated either overnight at 4°C or for 2 h at RT. The nerves were washed at least three times for 10 min each in the same buffer, followed by secondary antibody incubation for 2 h at RT. The secondary antibodies used were anti-rabbit STAR580 (Abberior, cat. ST580-1002, 1:200) and anti-mouse STAR635P (Abberior, cat. 2-0032-052-6, 1:200). For 3-color staining, the nerves were first labeled with anti-Palm2-p22 and its related secondary antibody STAR580, and subsequently labeled with primary antibodies against neurofilament H (Synaptic Systems, cat. 171 106, 1:500) and peripherin (Synaptic Systems, cat. 424 004, 1:500). The related secondary antibodies used were anti-chicken STAR GREEN (Abberior cat. STGREEN-1005, 1:200) and anti-guinea pig STAR 635P (Abberior, cat. 2-0112-007-1, 1:200). After the final washing steps, the nerves were teased on a #1.5 coverslip and mounted in Mowiol supplemented with DABCO.

2-color confocal images were acquired on an Abberior Instruments Infinity line equipped with 660 nm and 775 nm STED lasers. The following excitation lasers and detection windows were used to image STAR580 and STAR635, respectively: 561 nm 30%, 580 – 630 nm detection; 640 nm 2%, 650 – 760 nm detection. Pixel size was 80 nm in xy and 350 nm in z, dwell time: 7 µs, 1 line accumulation, pinhole size: 125 µm. 3-color confocal images were acquired on the Abberior Instruments Expert line microscope described below. The following excitation lasers, detection windows, and line accumulations were used to image STAR GREEN, STAR580 and STAR635, respectively: 488 nm 5 %, 505 – 540 nm detection, 2 line accumulations; 561 nm 10-30 %, 580 – 620 nm detection, 1 line accumulation; 640 nm 0.5 – 1 %, 650 – 725 nm detection, 1 line accumulation. Pixel size was 100 nm in xy and 350 nm in z or 120 x 120 x 120 nm for cross sections, dwell time: 10 µs, pinhole size: 100 µm.

### Western blotting

Tissue homogenates from adult mice (Fig. 1A) or littermate mouse pups of increasing postnatal age (Fig. 3D) were resolved by SDS-PAGE, transferred to PVDF membranes, and developed by Enhanced Chemiluminescence (ECL) following standard procedures. For Fig. S1H, nitrocellulose blots of brain homogenates from adult C57BL/6N and *Palm2^em1(IMPC)J^* mice were developed with the alkaline phosphatase color reaction. For Fig. 7E, yeast cell pellets were quick-extracted with the NaOH/mercaptoethanol/trichloroacetic-acid technique (*61*), equal quantities of total yeast proteins subjected to SDS-PAGE, and nitrocellulose blots developed by ECL with mouse anti-LexA-DBD (2-12, sc-7544, Santa Cruz Biotechnology).

For quantification of paralemmin isoforms in adult mouse brain, immunoblot signals of at least two different homogenates were calibrated on a dilution series of bacterially expressed, purified recombinant paralemmins on the same blot membrane. Blots were developed by ECL, signals collected on X-ray film, scanned, and quantified with Quantity One® (Bio-Rad).

### mEGFP-Palm2 CRISPR knock-in

To label endogenous Palm2 in cultured rat HPNs with mEGFP, the vector pORANGE (gift from H. MacGillavry, Addgene plasmid #131471; RRID:Addgene_131471) (*62*) was used as a template for cloning. The sequence of Palm2 from Rattus norvegicus (NC_005104.4, Gene ID: 103692368; on 15-Nov-2023, this record was replaced with Gene ID: 298024) was used to design guideRNAs (gRNAs) close to the 5’-end of Palm2 within exon 1. For this the CRISPR design tool of Benchling (https://benchling.com) based on scoring algorithms (*63, 64*) was applied. Synthetic DNA fragments encoding the designed gRNA sequences with overhangs for restriction digestion (Bbsl) were hybridized (Table S2, primers #1-2). After digestion of the vector, the gRNA-encoding insert was ligated into the pORANGE template. After ligation, transformation and purification, a second insert (mEGFP flanked by two inverse gRNA sequences) was generated via PCR (Table S2, primers #3-4). Another restriction digest was conducted with XbaI and Sall, and insert and vector were ligated. After bacterial transformation, the purified plasmid (GeneJET Endo-free Plasmid Maxiprep Kit, Thermo Fisher cat. K0861) was sequenced to ensure the correct design. Knock-in of mEGFP close to the 5’-end of Palm2 was verified by PCR and sequencing from whole genomic DNA (GenEluteTM Mammalian Genomic Miniprep Kits, Sigma-Aldrich G1N70-1KT) of primary rat hippocampal neurons, at least 7 days after electroporation with the plasmid. The layout of the resulting knock-in expression product is: M-linker1-mEGFP-linker2-Palm2, and its annotated sequence is given in Supplementary note 1.

### HPN culture preparation

All experiments were conducted in accordance with the Animal Welfare Act of the Federal Republic of Germany (Tierschutzgesetz der Bundesrepublik Deutschland, TierSchG) and the Animal Welfare Laboratory Animal Regulations (Tierschutz-Versuchstierverordnung), supervised by Animal Welfare officers of the Max Planck Institute for Medical Research (MPIMF) and of the Max Planck Institute for Multidisciplinary Sciences (MPINAT), conducted and documented according to the guidelines of the TierSchG (permit number assigned by the MPIMF:MPI/T-35/18 and MPI/T-36/18). For the procedures performed in this study, no specific authorization or notification was required. We have complied with all relevant ethical regulations for animal use.

Hippocampi were isolated from P0 to P2 post-natal WT Wistar rats (Janvier-Labs, Le Genest-Saint-Isle, France), C57BL/6N mice, or Palm2-KO mice of either sex and digested with 0.25% trypsin at 37°C. After 20 min, the reaction was stopped by adding 1× Dulbecco’s modified Eagle’s medium supplemented with 10 % heat-inactivated fetal bovine serum (FBS; Thermo Fisher Scientific cat. 12491015 and 10082147, respectively) and hippocampi were rinsed three times with Hanks’ solution (Thermo Fisher Scientific, cat. 10012011). Thereafter, they were mechanically dissociated by pipetting up and down in Neurobasal (NB) medium (Thermo Fisher Scientific, cat. 21103049) supplemented with 1 % GlutaMAX (Thermo Fisher Scientific, catalog no. 35050061), 1 % penicillin/streptomycin (Thermo Fisher Scientific, cat. 15070063), and 2 % B27 (Thermo Fisher Scientific, cat. 17504044) (from here on referred to as supplemented NB). Dissociated neurons were passed through a 40 μm cell strainer (Fisherbrand, cat. 22363547) before seeding them on glass coverslips precoated with poly-L-ornithine (0.1 mg/ml; Sigma-Aldrich, cat. P3655) and laminin (1 μg/ml; Corning, cat. 354232) with a total of 110,000 cells on ∅ 18 mm coverslips (12-well plate), or 55,000 cells on ∅ 12 mm glass coverslips (24-well plate). After 1-2 h, medium was replaced with fresh supplemented NB. After 24 h, 5 μM cytosine β-D-arabinofuranoside (AraC) was added to the cultures to inhibit cell division of glial cells and further incubated at 37°C and 5 % CO_2_ until use. Neurons that were not immediately used were placed directly after dissociation in cryotubes at a concentration of 3 to 4 million cells/ml (final media composition: 50 % supplemented NB, 40 % of heat-inactivated FBS, 10 % dimethyl sulfoxide) for freezing. Vials were placed into a freezing container and stored at −80°C overnight before being transferred into liquid nitrogen until further use. For thawing neurons, the frozen cryotubes were placed at 37°C until the cell solution was thawed. Then, cells were diluted 1:1 with supplemented NB and were transferred to fresh supplemented NB (end concentration 800,000 cells/ml). To achieve an expected density of ∼110,000 cells per coverslip, a total of 400,000 cells were plated on ∅ 18 mm coverslips (12-well plate). Finally, medium was changed to supplemented NB 1 h after seeding and neurons were further incubated at 37°C and 5 % CO_2_ until use.

### Electroporation of neurons

Freshly dissociated HPN were electroporated just before seeding at DIV 0 with the Neon Transfection System (Thermo Fisher Scientific, cat. MPK5000) and the Neon Transfection System 10 μl Kits (Thermo Fisher Scientific, cat. MPK1025) following the manufacturer’s instructions. Briefly, 150,000 neurons were washed once with PBS before adding 200 ng of plasmid and 10 μl of Buffer R. In the pipette tip, cells were electroporated with three pulses of 10 ms each at 1400 V and immediately plated on ∅ 18 mm glass coverslips (12-well plate) precoated as described above. Neurons were then incubated at 37°C and 5 % CO_2_ for 24 h, when the medium was changed to fresh supplemented NB medium with additional 5 μM AraC. Neurons were further incubated at the same conditions until use.

### YFP plasmids for transient overexpression

Mouse Palm2-coding sequences without N-terminal start codons but with C-terminal stop codons were amplified by RT-PCR from mouse brain single-stranded cDNA (Palm2: AEAEL…CVVM*, RefSeq: NP_766456, NM_172868). Inserts were cloned via the Gateway® system into a destination vector with an N-terminal YFP-encoding sequence (Invitrogen Vivid Colors® pcDNA6.2/N-YFP-DEST). The coding sequences of all recombinant plasmids were confirmed to be mutation-free by sequencing. The destination plasmid without paralemmin insert was used to express YFP as a control.

### Quantification of overexpression levels

To quantify overexpression levels, neurons were electroporated with YFP-Palm2 plasmid at DIV 0 and fixed in PFA at DIV 19. On the same coverslip, endogenous and recombinant Palm2 were immunostained with Palm2-1A8 antibody (1:100 dilution) and imaged using confocal microscopy. Local intensities of endogenous and recombinant Palm2, distinguished by the presence of YFP fluorescence only in transfected cells, were analyzed in FIJI by manually drawing a 6 µm-long line profile in the proximal and middle axon. Values were normalized to the average of each experimental round obtained in the proximal region of untransfected cells.

### Immunostaining of cultured neurons for confocal and STED imaging

Primary hippocampal neurons were briefly rinsed once with PBS and fixed in 4 % PFA in PBS for 20 min at room temperature, or 100 % methanol, or acetone for 10 min at -20°C before immunostaining. Only after PFA fixation, samples were quenched in quenching buffer (PBS, 100 mM glycine, and 100 mM ammonium chloride) and permeabilized for 5 min in 0.1 % Triton X-100 in PBS. Samples were then blocked with 1 % BSA in PBS for 1 hour and incubated with primary antibodies for 1 hour at room temperature in a wet and dark chamber. Samples were rinsed five times with PBS and secondary antibodies were added under the same conditions. Finally, samples were embedded in Mowiol 4-88 mounting medium (Merck, Sigma-Aldrich, cat. 81381) supplemented with 2.5 % w/w DABCO33-LV (Merck, Sigma-Aldrich, cat. 290734), according to the CSH protocol (https://cshprotocols.cshlp.org/content/2006/1/pdb.rec10255).

The following primary antibodies were used in addition to the Palm2 antibodies described above: βII-spectrin mouse (BD bioscience, cat. 612563, 1:400), ankyrin G guinea-pig (Synaptic Systems, cat. 386 005, 1:400), ankyrin G rabbit (Synaptic Systems, cat. 386 003, 1:400), ankyrin G guinea pig (Synaptic Systems, catalog no. 386005, 1:400), βIV-spectrin (a kind gift of Maren Engelhardt, 1:1000), Kv1.2 clone K14/16 (Neuromab, cat. 73-008, 1:10). The following secondary antibodies, nanobodies, and phalloidin conjugates were used at 1:100 dilution unless otherwise indicated: goat anti-mouse STAR635 (Abberior, cat. ST635P-1001), goat anti-rabbit STAR635P (Abberior, cat. ST635P-1002), goat anti-mouse STAR580 (Abberior ST580-1001), goat anti-rabbit STAR580 (Abberior, cat. ST580-1002), goat anti-guinea pig Alexa Fluor 488 (Thermo Fisher, cat. A-11073), FluoTag X-4 anti-GFP (Nanotag Biotechnologies, cat. N0304-Ab635P-S, 1:200), phalloidin-STAR635 (Abberior, cat. 2-0205-002-5).

### Confocal and STED imaging

Confocal and STED images were performed on an Abberior Instruments Expert Line Microscope (Abberior Instruments GmbH, Germany) built on a motorized inverted IX83 microscope (Olympus, Tokyo, Japan) and equipped with pulsed STED lines at 775 nm and 595 nm, RESOLFT lines at 488 nm and 405 nm, excitation lasers at 355 nm, 405 nm, 485 nm, 580 nm, and 640 nm, and spectral detection. Spectral detection was performed with avalanche photodiodes (APD) and detection windows were set to 650-725 nm, 600-630 nm, 505-540 nm, and 420-475 nm to detect STAR635P, STAR580, Alexa Fluor 488/YFP/eGFP and Alexa Fluor 405, respectively. Confocal images were acquired either with a 20x/0.4 NA oil immersion lens or with a 100x/1.4 NA oil immersion lens. The STED donut was generated with spatial light modulators (SLMs). STED images were acquired with the 100x/1.4 NA lens with a pixel size of 30 nm. Laser powers and dwell times were adjusted for the different experiments but kept consistent for the different conditions within the same experiment.

### Image processing and analysis

Acquired images (except Fig. 1B,C,G,H, S1A-C,F,G, see above) were visualized by Imspector (Abberior Instruments GmbH) and processed by FIJI ImageJ 1.52p (https://fiji.sc). Axonal regions were classified as described previously (*32*). Briefly, proximal region (=AIS, defined as ankG IF-positive), middle (up to 40 μm after the AIS), and distal (further than 40 μm after the end of the AIS).

Local intensities were obtained from the confocal images drawing manually line profiles of 6 μm along the axons in FIJI, sometimes including the same regions used for the correlation analysis. Values were normalized to the average of each experimental round of the respective control of (either nontransfected cells or YFP-transfected neurons) in the proximal regions. Only one proximal region, one middle region, and one distal region were measured per axon.

AC and CC analyses were performed as described previously (*32*). Briefly, regions of interest 1.5 to 2 μm long were manually selected and examined for periodic patterns using the function “xcorr2” of MATLAB. Amplitudes shown were calculated as the difference between the value at 190 nm and the average of the values at 95 and 285 nm, where the first two valleys are expected, from the AC curves.

Sholl analysis was performed on confocal images of neurons at DIV 3-4 with the FIJI tool “Sholl Analysis (From Image)”. Briefly, structures of the image that do not belong to the identified neuron were manually removed. After thresholding the image, all structures smaller than 0.5 μm were removed using “analyze particles”. Thereafter, a median filter with radius 1 was applied and the images were skeletonized. Next, contour of somata were manually drawn and their centroids were calculated to set the center of the concentric shells, spaced 1 μm apart. The last concentric shell did not intersect with any neuronal structure.

AIS length was measured as described in (*65*). A segmented line was then drawn from the soma to the end of the axon in the image, using the width of the axon’s widest part. Line profiles were smoothed using a 3 µm rolling average centered on the pixel of interest, then normalized by scaling the smoothed intensity values to range from 0 (minimum) to 1 (maximum). The start and end positions of the AIS were identified where these normalized values first dropped to 0.33 (the threshold) on either side of the maximum, even if they rose above the threshold again further along the profile. The AIS length was calculated by measuring the distance between the start and end positions.

Brightness was adjusted uniformly throughout the images for better display. No further image processing was performed and images are displayed as raw data, unless stated in the figure caption.

### Electrophysiological analysis of hippocampal neurons

Primary hippocampal neurons (DIV 20–26) derived from C57BL/6N WT or Palm2-KO mice were recorded using whole-cell patch clamp in either current-or voltage-clamp configuration. The following internal solutions were used for each configuration. For current-clamp recordings: 125 mM K-gluconate, 20 mM KCl, 10 mM HEPES, 0.5 mM EGTA, 4 mM MgATP, 0.3 mM NaGTP, and 10 mM Na-phosphocreatine (osmolarity 312 mOsm, pH 7.2 adjusted with KOH). For voltage-clamp recordings: 125 mM Cs-gluconate, 20 mM KCl, 4 mM MgATP, 10 mM Na-phosphocreatine, 0.3 mM NaGTP, 0.5 mM EGTA, 2 mM QX-314, and 10 mM HEPES (312 mOsm, pH 7.2). The extracellular solution used for both configurations was continuously oxygenated with 95% O_2_ and 5% CO_2_ and contained: 125 mM NaCl, 2.5 mM KCl, 25 mM NaHCO_3_, 0.4 mM ascorbic acid, 3 mM myo-inositol, 2 mM Na-pyruvate, 1.25 mM NaH_2_PO_4_, 2 mM CaCl_2_, 1 mM MgCl_2_, and 25 mM D(+)-glucose (315 mOsm, pH 7.4).

For electrophysiological recordings, a 12-mm glass coverslip (∅ 12 mm, from a 24-well plate) containing approximately 55,000 neurons was placed in an RC-27 recording chamber (Warner Instruments) and mounted on a BX51 upright microscope (Olympus) equipped with DIC and fluorescence capabilities. Bath temperature was maintained at 26 ± 1°C using a dual-channel TC-344B temperature controller (Warner Instruments). Neurons were visually identified and patched under DIC optics using borosilicate glass pipettes with a resistance of 3–4 MΩ (WPI), pulled with a PC-100 puller (Narishige, Japan). Signals were amplified using a Multiclamp 700B amplifier (Molecular Devices) and acquired with Clampe× 10.1 software via a Digidata 1440A digitizer (Molecular Devices).

For current-clamp recordings, the membrane potential was held at approximately −70 mV by injecting a constant holding current; only cells requiring less than 50 pA to maintain this potential were included. To determine the rheobase — defined as the minimum current required to trigger an action potential (AP) — neurons were depolarized from resting potential by successive injections of positive current in 1-pA steps (500 ms duration). AP properties (height and half-width) were extracted from spikes evoked during rheobase measurements. Input resistance and input/output relationships were assessed using stepwise current injections (500 ms, 25-pA steps ranging from −200 to +400 pA).

For voltage-clamp recordings, only cells with a series resistance below 10 MΩ were included; series resistance was not electronically compensated. Miniature excitatory postsynaptic currents (mEPSCs) were recorded in the presence of 0.5 μM TTX and identified as downward deflections in the current trace. Current-and voltage-clamp recordings were analyzed using Clampfit 10.1 or custom-written macros in IgorPro 6.11 (WaveMetrics).

### Yeast-2-hybrid assays (Y2H)

All Y2H assays were performed at Hybrigenics (Evry, France). A human fetal brain cDNA library in the prey plasmid pP6 (Hybrigenics) was screened with the bait construct of mouse Palm1ΔEx8 (aa 1-326; MEVLA…DTQDL; full-length except for the last 13 codons) in plasmid pB27 (layout of the bait fusion protein: N-LexA-Palm1ΔEx8-C). Palm1ΔEx8 was preferred as the screening bait over Palm2 for two reasons: (i) It has no autoactivation in the Y2H test even at 0 mM 3AT, whereas full-length Palm2 displays autoactivation at 0-1 mM 3AT, and therefore avoids unspecific background in the screen; (ii) It lacks spectrin binding to homology region 2 which is encoded by Palm1 exon 8 and constitutively expressed in Palm2, and therefore avoids a high specific background of β-spectrin isoforms. Screening at 0 mM 3AT yielded two USP7 prey clones with inserts of aa 872-1066 (QPKK…FEPQ) and three prey clones with inserts of aa 891-1074 (RRSF…SHPR).

For subsequent pairwise tests, the mouse Usp7 sequence PKKL…SHPR (aa 874-1075; RefSeq NP_001003918; collinear to the composite human bait sequences derived from the library screen) was cloned into prey plasmid pP7 (Hybrigenics). In addition to the bait N-LexA-Palm1ΔEx8-C in pB27, as used in the screen, paralemmin structural variants and isoforms were cloned into the bait vector pFBL23. pFBL23 expresses the bait fusion proteins with the paralemmin sequences at the N-terminus, followed by the LexA DNA-binding domain (N-Palm1ΔEx8-LexA-C). See (*32*) for further details of bait vector design. The following Palm1 bait sequences in pFBL23, to be tested for interaction with Usp7, were described before (*32*): Palm1ΔEx8-LexA, Palm1, Palm1-E5, Palm1(W54A) (all full-length, except for the last 13 codons) and Palm1(1–263). Additionally, mouse Palm2 (MAEAE…DPAPG; full-length except for the last 13 codons) was cloned into pFBL23 in the same sequence context. Pairwise Y2H assays were performed as described in detail previously (*32*) and included controls with empty bait and prey plasmids to test for autoactivation by the respective preys and baits.

The prey series of β-spectrin/actinin isoforms, modeled on the βII-spectrin sequence “construct G” (aa 185-411; MKTA…LALR), each encompassing the truncated CH2 domain, the Palm1-binding site, and the first spectrin repeat SR1, was described and tested for interaction with Palm1 before (*32*). As Palm2 baits for testing interactions with β-spectrin/actinin isoforms, mouse and human partial sequences were cloned into pFBL23, again exactly as described in (*32*) for Palm1. As the N-terminal two thirds of Palm1 (construct Palm1(1–263)) retain the βII-spectrin binding activity (see Fig. 6G of (*32*)), the mPalm2 and hPalm2 baits were designed analogously: mPalm2(1–266) (aa 1-266; MAEA…ITEE; RefSeq: NP_766456, NM_172868) and hPalm2(1–301) (aa 1-301; MAEA…SAAG; RefSeq: NP_443749, NM_053016). These partial Palm2 bait sequences were preferred over the full-length sequences to reduce the autoactivation background which Palm2, unlike Palm1, displays in the Y2H system. Residual autoactivation background of hPalm2(1–301) can be seen in Fig.7B as traces of yeast growth at 0 mM 3AT with the preys βIII-and βV-spectrin, and actinin-2. In Fig. 7C-E, these truncated baits are simply referred to as mPalm2 and hPalm2 for brevity. In separate experiments with full-length mPalm2, above-background interactions with βI-, βII-and βIV-spectrins and also weakly with Actn1 and Actn4 were confirmed. 3-Amino-1,2,4-triazol (3AT) is a competitive inhibitor in the histidine biosynthesis pathway that enhances the stringency of growth selection, and a concentration series of 3AT gives a measure of the relative strength of bait-prey interaction.

### Statistical analysis and preparation of figures

Statistical tests and the plotting of graphs were performed with Origin 2019 and are described in each caption. Correlation coefficients were interpreted as suggested in (*66*): 0.0-0.3 no correlation, 0.3-0.5 low correlation, 0.5-0.7 moderate correlation, 0.7-1 high correlation. Statistical differences are indicated as follows: * = p<0.05; ** = p<0.005, *** = p<0.0005.

All figures were assembled in Adobe Illustrator 2022.

## Acknowledgements

We thank Prof. Stefan W. Hell and Prof. Nils Brose for supporting this project. We acknowledge Kathrin Kusch and Christiane Senger-Freitag (Institute for Auditory Neurosciences, Göttingen) for re-cloning the full-length Palm2 insert into pFBL23, Astrid Zeuch for immunoblot analysis of yeast extracts, and Petra Tafelmeyer (Hybrigenics) for expert Y2H support. We thank Francisco Balzarotti for developing the MATLAB script to measure the periodicity; Gabriel Tulcan for support in AIS length analysis. We are grateful to Katrin Willig for access to laboratory spaces; Alena Fischer, Jana Kress, Birgit Koch, Paula Breuer for support with neuronal cultures; and Valerie Dürr and Daniel Bollack for support with sample preparation. We thank Torben Ruhwedel and Wiebke Möbius for support with animal perfusion, Sua Jeong and JeongSeop Rhee for guinea-pig tissue, the staffs of the animal facility and the genotyping facility of the MPI for Multidisciplinary Sciences (City Campus) for Palm2-KO mouse generation, maintenance, breeding and genotyping, and Hauke Werner for scientific input on sciatic nerve experiments. This research was funded by the Deutsche Forschungsgemeinschaft (DFG, SFB1286/A07 to E.D.; grant KI 324/14 to M.W.K), and the Swedish Research Council (Vetenskapsradet 2003-3398) to M.W.K..

## Author contributions

V.M.P. performed and analyzed all cell culture and CNS NoR imaging experiments, and prepared the figures, supervised by E.D.; N.G.M. generated YFP expression plasmids and performed exploratory IF & overexpression experiments, noting Palm2 to be enriched at the proximal axons, and G.H. generated & validated Palm2 antibodies and performed immunoblotting of adult mouse organs, both supervised by M.W.K.; C.A. performed and analyzed electrophysiology experiments; J.H. designed the CRISPR/Cas9 construct; S.K. performed CRISPR experiments supervised by J.H. and V.M.P; L.W. and H.M. performed immunohistology of adult tissues and immunoblotting of KO mouse brain in interaction with M.W.K.. M.W.K. designed and supervised the Y2H analyses, and provided the Palm2-KO mice; E.D. prepared and imaged sciatic nerve samples. V.M.P., E.D. and M.W.K. conceived the experiments and conceptualized the project, and wrote the manuscript with input from all authors.

## Animal Ethics declaration

All regulations given in §4 Animal Welfare Law of the Federal Republic of Germany (§4 TierSchG, Tierschutzgesetz der Bundesrepublik Deutschland) and the Animal Welfare Laboratory Animal Regulations (Tierschutz-Versuchstierverordnung) were followed and supervised by Animal Welfare officers of the Max Planck Institute for Medical Research (MPIMF) and of the Max Planck Institute for Multidisciplinary Sciences (MPINAT), conducted and documented according to the guidelines of the TierSchG (permit number assigned by the MPIMF: MPI/T-35/18 and MPI/T-36/18). No specific authorization or notification was required for the breeding of animals, or for the procedures performed in this study. We have complied with all relevant ethical regulations for animal use.

## Supplemental information

**Document S1:** Figures S1-S4, Supplementary note 1, Supplementary tables 1 and 2.

**Data S1**: Excel file containing the p-values for all statistical tests performed, related to all figures.

## Data and materials availability

All data needed to evaluate the conclusions in the paper are present in the paper and/or the Supplementary Materials. The C57BL/6NJ-*Palm2^em1(IMPC)J^*/Mmjax mouse line is available at the Mutant Mouse Resources & Research Centers (MMRRC) with the stock number 42398-JAX.

Palm2 plasmids and antibodies are available from MWK, except the plasmid for the CRISPR/Cas9 Palm2 tagging which is available from ED.

