## Supplementary Information for "Paralemmin-2 is a membrane-anchored cytoskeletal constituent of Axon Initial Segments and nodes of Ranvier"

### Supplementary Figures

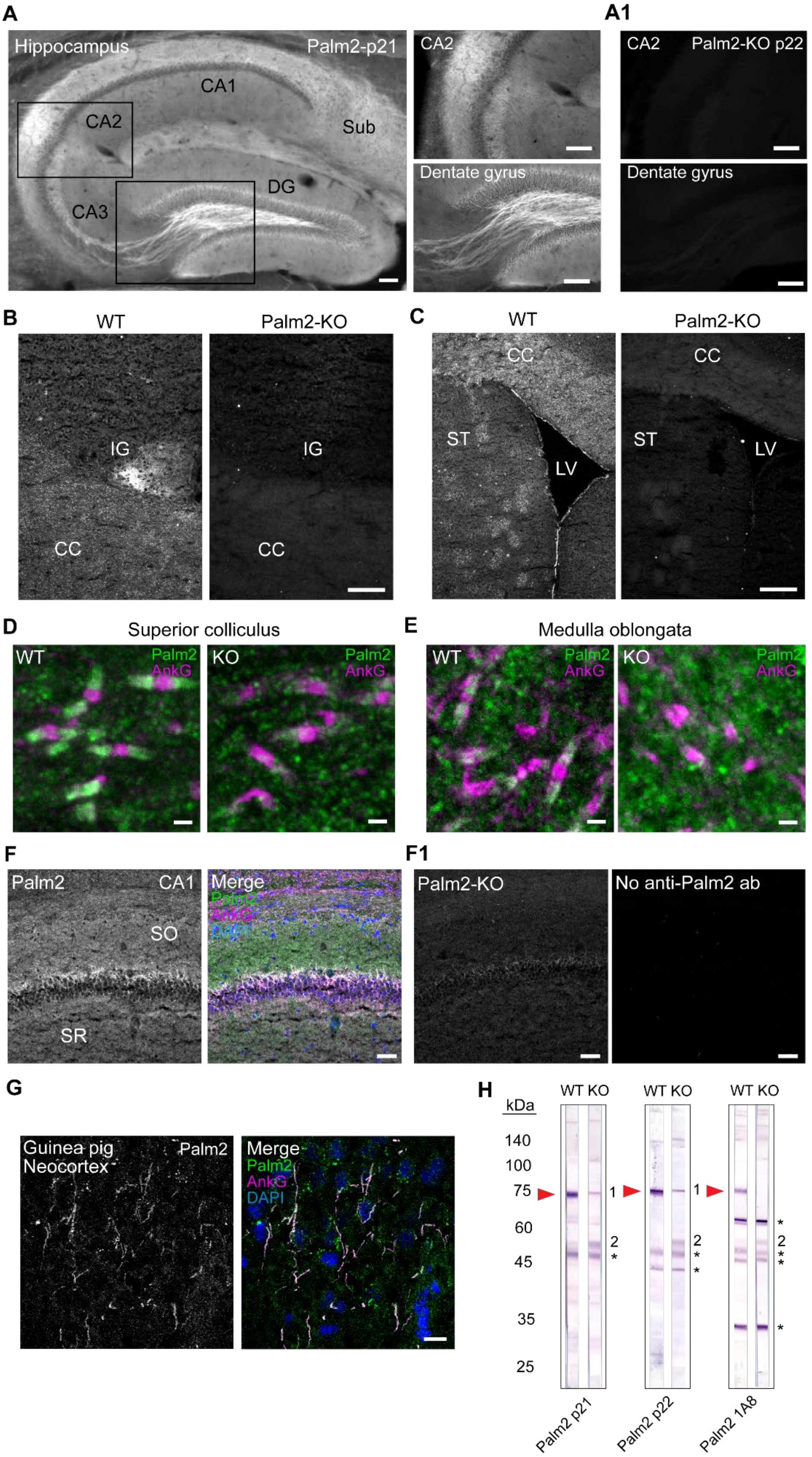

**Figure S1: Palm2 localizations and specificity controls (independent antibodies and KO tissues).** (A) In hippocampus, staining of axons and neuropil for Palm2 is positive with anti-p21, like with anti-p22 as shown in Fig. 1B. Note that even subtle features like the enhanced staining of the CA2 stratum oriens and mossy fibers are reproducible with both antibodies (close-ups on the right). Tissue PFA-fixed, Triton-permeabilized, heat-treated. Scale bars: 100 µm. (A1) Signal is markedly reduced in the CA2 and dentate gyrus of Palm2-KO tissue stained with anti-p22 staining. Compare to WT tissue in Fig. 1B1 and 1B2. Tissue PFA-fixed, Triton-permeabilized, heat-treated. Scale bars: 100 µm. (B) Field of view showing the corpus callosum (CC)/neocortex boundary encompassing the indusium griseum (IG). Both the NoR staining in the corpus callosum and the finely dispersed staining in the indusium griseum are abolished in the Palm2-KO tissue. Anti-p22 on fresh-frozen, acetone-postfixed tissue. Scale bar: 100 µm. (C) Field of view showing the corpus callosum (CC, top), the striatum (ST, bottom left), and the lateral ventricle (LV, bottom right). Palm2-IF of NoR in the corpus callosum and in cross-sectioned myelinated fiber bundles of the striatum, as well as in the ventricle lining, is nearly abolished in the Palm2-KO tissue (anti-p22 on fresh-frozen acetone-postfixed tissue). Scale bar: 100 µm. (D, E) At NoR in the brainstem (superior colliculus, medulla oblongata), Palm2-IF (anti-1A8) localizes to paranodes and is nearly abolished in KO mouse tissue (PFA-perfused mice). Scale bars: 1 µm. (F) Palm2 co-enrichment with ankG at AIS in the stratum pyramidale of the hippocampal CA1 area, along with fainter diffuse staining in the stratum oriens (SO) above and stratum radiatum (SR) below, is seen in fresh-frozen, acetone-postfixed specimens also with anti-p22 staining. Scale bar: 20 µm. (F1) Palm2-IF is markedly reduced in Palm2-KO mouse tissue, and absent in the reagent control without anti-Palm2. Scale bar: 20 µm. (G) Palm2 is AIS-enriched also in guinea-pig neocortex, as in mouse brain. Anti-p21 on fresh-frozen/acetone-postfixed specimen. Scale bar: 20 µm. (H) Immunoblots of WT and KO mouse neocortex homogenates, developed with three independent Palm2 antibodies as indicated on the bottom. Red arrowheads next to the WT lanes indicate the specific Palm2 band of 72 kDa. Next to the KO lanes, “1” and “2” mark aberrant bands newly appearing or intensified in the KO tissue, whereas asterisks indicate unspecific bands depending on the respective antibody and present in both WT and KO samples. Aberrant band 1 in the KO is slightly but consistently higher than the regular 72 kDa band of the WT. See Table S1 for experimental details.

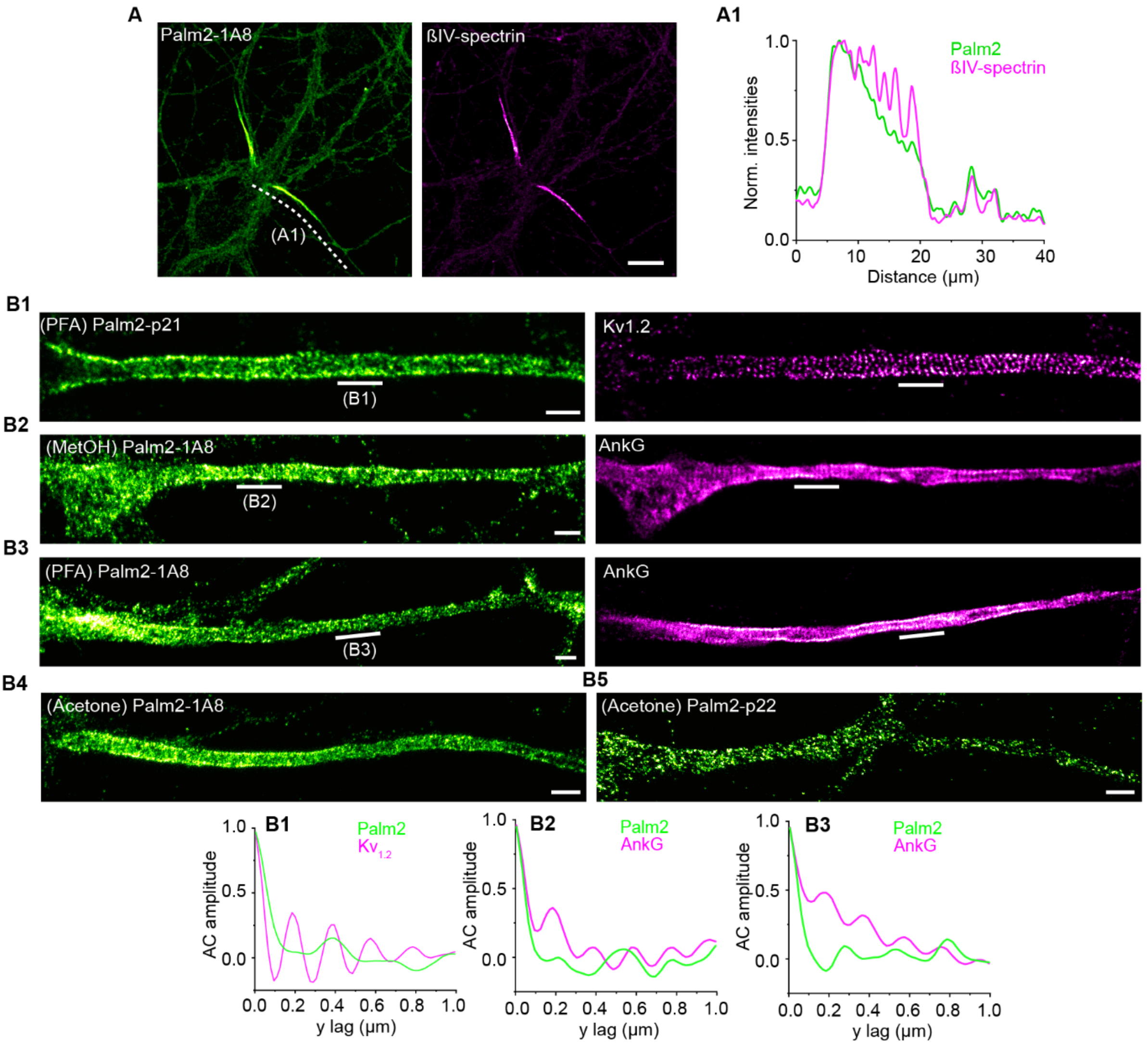


**Figure S2: Endogenous Palm2 is non-periodic regardless of the antibody and fixation conditions.**

(A) Representative confocal image of a rat HPN at DIV 19 immunolabeled against endogenous Palm2 (1A8 antibody) and βIV-spectrin. Scale bar: 10 µm. (A1) Smoothed and normalized fluorescence intensities of the proteins along the axon highlighted by the dashed line in (A). (B1-5) Representative STED images of mature HPN immunolabeled with different Palm2 antibodies upon different fixation methods, as indicated in each panel in green. On the right in magenta: co-staining with selected AIS markers that exhibit a periodic organization. Scale bars: 1 µm. On the bottom: AC analysis of the AIS regions indicated by the lines show periodic patterns for $Kv_{1.2}$ and ankG, but not for Palm2.

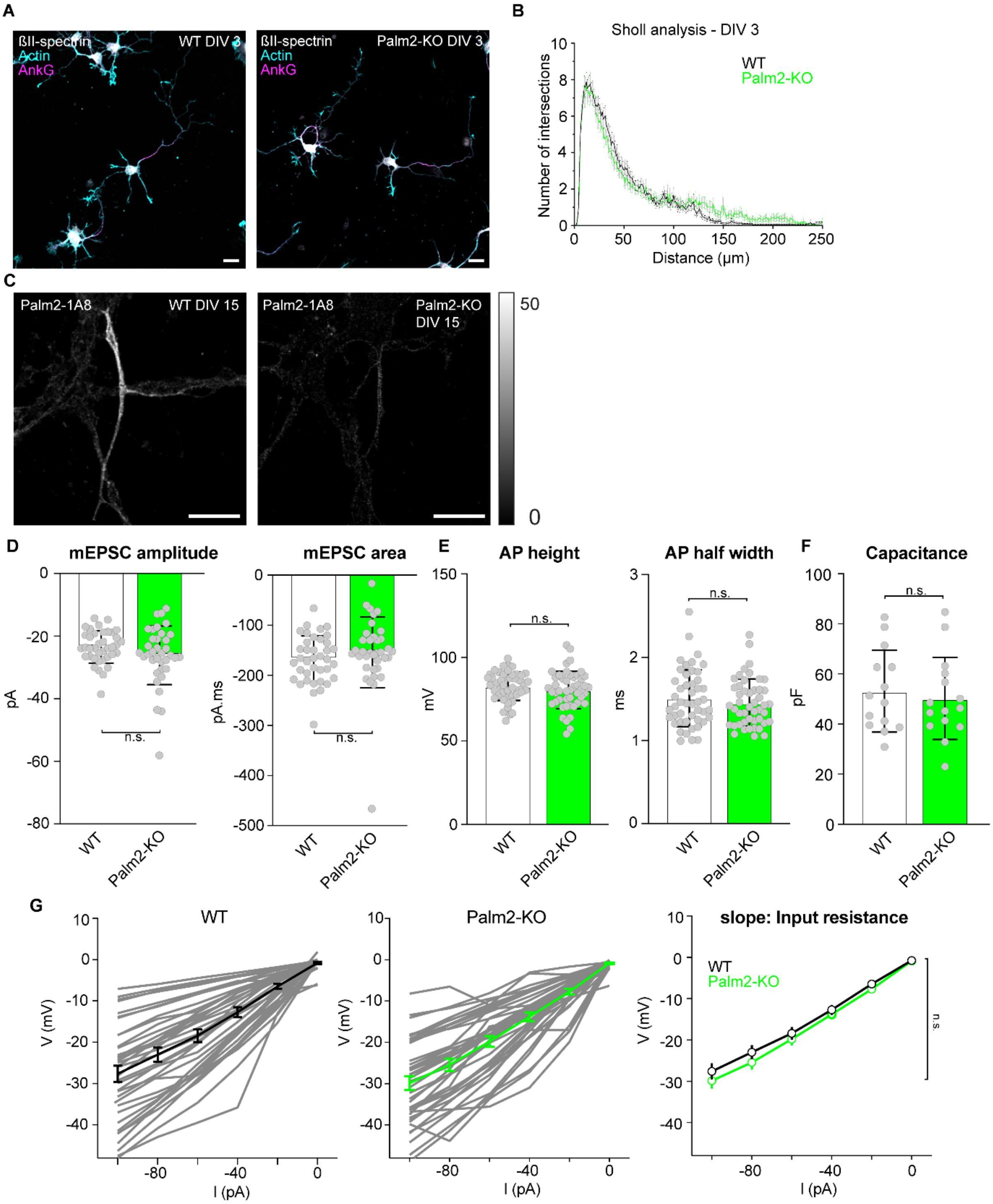

A
ßII-spectrin
Actin
AnkG
WT DIV 3
Palm2-KO DIV 3
B
Sholl analysis - DIV 3
WT
Palm2-KO
Number of intersections
Distance (µm)
C
Palm2-1A8
WT DIV 15
Palm2-KO DIV 15
D
mEPSC amplitude
mEPSC area
pA
pA.ms
E
AP height
AP half width
mV
ms
F
Capacitance
pF
n.s.
WT
Palm2-KO
G
WT
Palm2-KO
slope: Input resistance
V (mV)
I (pA)

**Figure S3: Phenotypic features of Palm2-KO neurons.** (A) Immature Palm2-KO neurons show a morphology similar to WT. Representative confocal images of WT and Palm2-KO neurons at DIV 3. Scale bar: 20 µm. (B) Sholl analysis of Palm2-KO neurons at DIV 3 showed a similar branching than WT neurons, but longer processes. Number of neurons analyzed WT/KO: 31/25. N = 3. Area under curve (AUC) 0-120 µm WT/KO mean ± S.D.: 372,9 ± 120,7 / 349,1 ± 88,4, p-value = 0,4; AUC 120-200 µm WT/KO mean ± S.D.: 15,6 ± 25,2 / 54,1 ± 64,3, p-value: 0,008. (C) Representative confocal image of an AIS of a (left) WT and (right) Palm2-KO mature neuron (DIV 15) immunolabelled against Palm2 (1A8 antibody), showing a low residual Palm2-IF in the AIS Palm2-KO neurons. Scale bars: 10 µm. (D) Unaffected electrophysiological parameters of mEPSCs, (E) of action potentials, (F) capacitance, and (G) input resistance from WT and Palm2-KO neurons. Same dataset as Fig. 4. Statistical analysis for all histograms: Paired Sample T-Test.

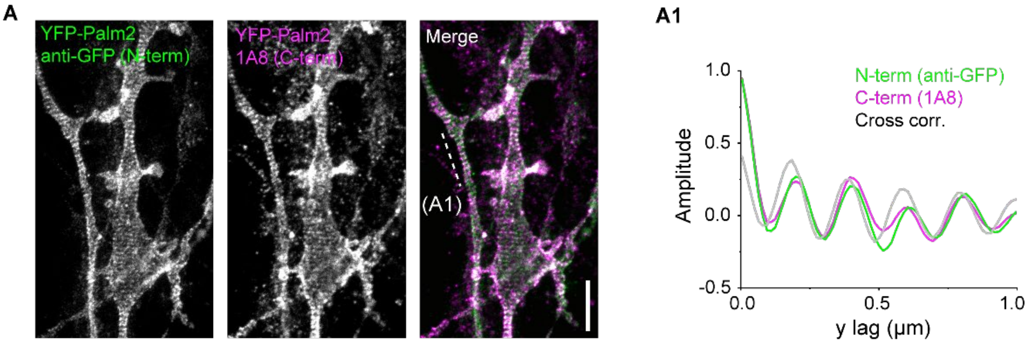


**Figure S4: Overexpressed YFP-Palm2 is periodic, regardless of N- or C-terminal labeling.** (A) Representative STED images of an axon and a dendrite of the same neuron overexpressing YFP-Palm2, immunolabeled against both its N-terminus (anti-GFP nanobody) and the C-terminus (Palm2-1A8). Scale bar: 2 µm. (A1) AC and cross correlation amplitudes obtained from the two different stainings along the axon, as indicated by the dashed line, show an overlapping periodic pattern.

### Supplementary note 1

DNA and corresponding amino acid sequences of mEGFP-Palm2 CRISPR/Cas9 knock-in

Color code: ATG + 1bp exon 1 Palm2, gRNA, Linker, mEGFP, Linker, 3bp inverse gRNA, Palm2

ATGGCCATCCTGGAGAAGGGCGCTAGCATGGTGAGCAAGGGCGAGGAGCTGTTCACCGGGGT
GGTGCCCATCCTGGTCGAGCTGGACGGCGACGTAAACGGCCACAAGTTCAGCGTGTCCGGCGA
GGGCGAGGGCGATGCCACCTACGGCAAGCTGACCCTGAAGTTCATCTGCACCACCGGCAAGCT
GCCCGTGCCCTGGCCCACCCTCGTGACCACCCTGACCTACGGCGTGCAGTGCTTCAGCCGCTA
CCCCGACCACATGAAGCAGCACGACTTCTTCAAGTCCGCCATGCCCGAAGGCTACGTCCAGGA
GCGCACCATCTTCTTCAAGGACGACGGCAACTACAAGACCCGCGCCGAGGTGAAGTTCGAGGG
CGACACCCTGGTGAACCGCATCGAGCTGAAGGGCATCGACTTCAAGGAGGACGGCAACATCCT
GGGGCACAAGCTGGAGTACAACTACAACAGCCACAACGTCTATATCATGGCCGACAAGCAGAAG
AACGGCATCAAGGTGAACTTCAAGATCCGCCACAACATCGAGGACGGCAGCGTGCAGCTCGCC
GACCACTACCAGCAGAACACCCCCATCGGCGACGGCCCCGTGCTGCTGCCCGACAACCACTAC
CTGAGCACCCAGTCCAAGCTGAGCAAAGACCCCAACGAGAAGCGCGATCACATGGTCCTGCTG
GAGTTCGTGACCGCCGCCGGGATCACTCTCGGCATGGACGAGCTGTACAAGGGCTCGAGCCCA
TCAACAAGTTTGTACAAAAAAGCAGGCTCCGCGGCCGCCCCCTTCACCGCCTCTGCAGAGGCG
GAATTGCACAAGGAGAGGCTGCAAGCCATAGCAGAAAAAAGAAAGAGGCAGACGGAAATAGAAG
GGAAGAGACAACAGCTTGACGAGCAGGTGCTGCTGCTCCAACATTCCAAGTCCAAAGTGCTTCG
GGAAAAATGGCTGCTGCAGGGTGTACCGGCGGGAACAGCGGAGGAGGAGGAAGCCAGACGCC
GACAGTCTGAGGAGGATGAATTCAAAGTCAAGCAGCTTGAAGATAACATTCAGAGGCTGGAGCA
GGAAATACAAGCGCTCGAAAGCGAAGAGTCCCAGATATCGGCCAAAGAGCAGATCATCCTCGAG
AAACTGAAGGAGACAGAGAAATCCTTCAAGGACTTGCAGAAGAGTTTCTCCACTGCTGATGGAG

CTATATACGCCATGGAAATTAATGTGGAGAAAGACAAACAAACAGGAGAGACCAAGATTCTCTCT
GCATCCACCATTGGCCCAGAGGGGGTCCATCAGAGAGGAGTCAAAGTCTATGATGATGGTACCA
AAGTAGTGTATGAGGTGCACTCAGGAGGCACCGTGGTAGAGAACGGAGTCCACAAACTAAGCG
CAAAGGATGTGGAAGAGCTAATTCAGAAGGCTGGACAATCGAGCTTCAGAGGACACATGTCAGA
AAGAACTGTTGTTGCAGATGGGAGCCTAGGCCATCCCAAGGAACACATGCTCTGCAAAGAGGCC
AAGTTAGAAATGGTGCAAAAATCCAGGAAAGATCAGTCTTCGGGAAACCCCGGGCACCAAGCCC
AACCCCCCAGCACAGAGGGTCCAGAGGTCAACCTGGATCAACCGGTCACCATGATTTTTATGGG
CTACCAAAACATCGAGGATGAAGAGGAGACTAAGAAGGTGCTAGGCTATGACGAAACCATCAAG
GCCGAATTAGTGCTGATTGACGAAGACGATGAAAAGTCGCTGAGGGAGAAGACAGTGACGGATG
TGTCCACGATCGATGGGAATGCAGCTGAACTGGTGTCCGGCAGGCCCATGTCGGACACCACAG
AACCCTCATCCCCAGAAGACAAGGAAGAGAGCCTGGCCACAGACCCGGCCCCAGGTACCCAAA
AGAAAAAGCGCTGTCAATGCTGTGTTGTCATGTGA

AA sequence mEGFP, rat Palm2

MAILEKGASMVSKGEELFTGVVPILVELDGDVNGHKFSVSGEGEGDATYGKLTLKFICTTGKLPVPWP
TLVTTLTYGVQCFSRYPDHMKQHDFFKSAMPEGYVQERTIFFKDDGNYKTRAEVKFEGDTLVNRIEL
KGIDFKEDGNILGHKLEYNYNSHNVYIMADKQKNGIKVNFKIRHNIEDGSVQLADHYQQNTPIGDGPVL
LPDNHYLSTQSKLSKDPNEKRDHMVLLEFVTAAGITLGMDELYKGSSPSTSLYKKAGSAAAPFTASAE
AELHKERLQAIAEKRKRQTEIEGKRQQLDEQVLLLQHSKSKVLREK**W**LLQGVPAGTAEEEEARRRQS
EEDEFKVKQLEDNIQRLEQEIQALESEESQISAKEQIILEKLKETEKSFKDLQKSFSTADGAIYAMEINVE
KDKQTGETKILSASTIGPEGVHQRGVKVYDDGTKVVYEVHSGGTVVENGVHKLSAKDVEELIQKAGQ
SSFRGHMSERTVVADGSLGHPKEHMLCKEAKLEMVQKSRKDQSSGNPGHQAQPPSTEGPEVNLD
QPVT**MIF**MGYQNIEDEEET**KK**VLGYDETIKA**E**LVLIDEDDEKSLREKTVTDVSTIDGNAAELVSGRPMS
DTTEPSSPEDKEESLATDPAPGTQKKKRCQCCVVM

AA sequence features:

KSKVLREK**W**LL: Paralemmin motif with W50 in homology region 1

AIYAMEINVEKDKQTGETKILSASTIGPEGVHQRGVKVYDDGTKVVYEVHSGGTVVENGVHKLSAKDV
EELIQKAG: homology region 2 – Paralemmin core domain

GQSSFRGHMSERTVVADGSLGHPKEHMLCKEAKLEMVQKSRKDQSSGNPGHQAQPPSTEGPEVN
LDQ: p21 sequence

PVT**MIF**MGYQNIEDEEET**KK**VLGYDETIKA**E**LVLIDEDDEKSL: homology region 3 with MIF (F279, equivalent of Palm1ΔEx8 F257), KK292/293 (equivalent of Palm1ΔEx8 KK270/271), and E304 (equivalent of Palm1ΔEx8 E282)

KSLREKTVTDVSTIDGNAAELVSGRPMSDTTEPSSPEDKEESLATDPAPGTQK: p22 sequence

KKKRCQCCVVM: CaaX motif

### Table S1

Experimental details related to Fig. 1 and Fig. S1. PFA = paraformaldehyde; AGR = antigen retrieval with heat treatment

| Figure | Animal | Age | Fixation/sample treatment | Palm2 antibody |
|---|---|---|---|---|
| 1B | WT | 66 weeks | PFA/AGR | p22 |
| 1C | WT | 76 weeks | Fresh-frozen/Acetone | p22 |
| 1D | WT | p67 | PFA/AGR | 1A8 |
| 1E | WT | 92 weeks | PFA/Methanol | p22 |
| 1F | WT | 92 weeks | PFA/Methanol | p22 |
| 1G | WT | 12 weeks | Fresh-frozen/Acetone | p21 |
| 1H | WT | 66 weeks | PFA/AGR | p21 |
| S1A | WT | 66 weeks | PFA/AGR | p21 |
| S1A1 | Palm2-KO | 66 weeks | PFA/AGR | p22 |
| S1B | WT / Palm2-KO | 76 weeks | Fresh-frozen/Acetone | p22 |
| S1C | WT / Palm2-KO | 76 weeks | Fresh-frozen/Acetone | p22 |
| S1D | WT / Palm2-KO | p67 | PFA/AGR | 1A8 |
| S1E | WT / Palm2-KO | p67 | PFA/AGR | 1A8 |
| S1F | WT / Palm2-KO | 76 weeks | Fresh-frozen/Acetone | p22 |
| S1G | Guinea pig WT | adult | Fresh-frozen/Acetone | p21 |

### Table S2

Oligonucleotides used for the generation of the mEGFP-CRISPR vector. Forw = forward; rev = reverse.

| No. | Primer name | Sequence 5´- 3´ | Function |
|---|---|---|---|
| **1** | ORANGE_Palm-KI-gRNA4-forw | CACCGCCCTTCTCCAGGATGGCAG | gRNA for tagging Palm2 |
| **2** | ORANGE_Palm-KI-gRNA4-rev | AAACCTGCCATCCTGGAGAAGGGC | gRNA for tagging Palm2 |
| **3** | PCR_gRNA4_Palm2 forw | CTGCAGACAAATGGCTCTAGAAGCTTC CTCTGCCATCCTGGAGAAGGGCGCTAG CATGGTGAGCAAGGGCGAG | Forw primer for PCR for amplifying insert mEGFP |
| **4** | PCR_gRNA4_Palm2 rev | ACGCGTCCTAGGATCCTCGAGTCGACA ATTGGCCCTTCTCCAGGATGGCAGAGG CGGTGAAGGGGGCGGCCGC | Rev primer for PCR for amplifying Insert mEGFP |

### Video S1

**Animation of human Palm2 structure.** The relevant sequences are color-coded as in Fig. 7F. Further highlighted are W50 (equivalent of Palm1 W54), F314 (equivalent of Palm1 F257), and E339 (equivalent of Palm1 E282). Created with PyMol 3.1.7.2.
